# Transferrin receptor 1 binds human parvovirus B19 VP1u to facilitate entry

**DOI:** 10.64898/2026.03.12.711364

**Authors:** Hyunwook Lee, Jan Bieri, Nicolas Ammann, Corinne Suter, Daniela Hunziker, Ajit K. Singh, Susan L. Hafenstein, Carlos Ros

**Affiliations:** The Hormel Institute, University of Minnesota, Austin, MN 55912, USA; Department of Chemistry, Biochemistry and Pharmaceutical Sciences, University of Bern, Switzerland; Graduate School for Cellular and Biomedical Sciences, Bern, Switzerland; University of Minnesota, Department of Biochemistry, Biophysics and Molecular Biology, Minneapolis MN 55455, USA; Department of Infectious Disease, Mayo Clinic, 200 First St. SW, Rochester, MN 55905 USA

**Keywords:** Parvovirus B19, B19V, erythroid progenitor cells, VP1u, transferrin receptor 1, TfR1, CD71, cryoEM structure, virus uptake, tropism

## Abstract

Human parvovirus B19 (B19V) exhibits a strict tropism for erythroid progenitor cells, which is governed by the VP1 unique region (VP1u). This region mediates cell-specific uptake by interacting with an unknown cellular receptor, termed VP1uR. Proximity labeling in permissive erythroid cells identified transferrin receptor 1 (TfR1/CD71) as the predominant membrane protein associated with VP1u. VP1u constructs colocalized with TfR1 at the cell surface of erythroid cells. Incubation with anti-TfR1 antibody OKT9 abolished binding and uptake of recombinant VP1u. While OKT9 efficiently inhibited B19V uptake and infection, it did not block virus binding to host cells. Direct binding assays confirmed interaction of VP1u to human TfR1. Using cryoEM we solved the 2.4 Å structure of the TfR1-VP1u complex, mapping the binding site and identifying the specific interactions. These findings establish TfR1 as the previously unknown receptor, VP1uR, required for B19V uptake.

## Introduction

Human parvovirus B19 (B19V), the prototype member of the genus *Erythroparvovirus*, is a common human pathogen associated with a range of clinical manifestations, including erythema infectiosum, transient aplastic crisis, chronic anemia in immunocompromised individuals, and severe fetal disease following intrauterine infection^1^. A defining feature of B19V biology is its exceptionally narrow cellular tropism: productive infection is restricted exclusively to erythroid progenitor cells (EPCs) in bone marrow and fetal liver^2,3^. This stringent restriction is a central determinant of viral pathogenesis.

Early studies identified the glycosphingolipid globoside as the cellular receptor for B19V^4^. However, globoside expression is not restricted to erythroid cells, and subsequent work demonstrated that globoside alone cannot account for the selective uptake and tropism of the virus^5^. Genetic and functional analyses showed that globoside is dispensable for viral uptake but essential at a post-entry step required for productive infection^6^. Recent work established globoside as a conditional receptor whose function is controlled by the local pH conditions^7^. At the mildly acidic airway mucosa, B19V binds globoside to mediate uptake and transcytosis across the respiratory epithelium^8^. In erythroid progenitor cells, globoside is dispensable for uptake but, upon endosomal acidification, it becomes essential for viral escape into the cytosol^9^. Thus, globoside acts as a conditional receptor supporting entry through the respiratory epithelium and endosomal escape in a strict pH-dependent manner.

B19V capsid is composed of VP1 and VP2. VP1 has an N-terminal extension of 227 amino acids, the so-called VP1 unique region (VP1u). Previous work demonstrated that B19V uptake is a highly selective process controlled by VP1u and restricted to permissive erythroid cells^10,11^. Subsequent mapping and mutational analyses localized a discrete receptor-binding domain (RBD) within the N-terminal region of VP1u that is necessary and sufficient to mediate erythroid-restricted internalization^12^. The unknown specific host receptor that is recognized by VP1u has been termed VP1uR^13^.

The functional properties of the VP1u RBD have been extensively characterized using heterologous display systems. Multivalent presentation of VP1u on different bacteriophage particles recapitulates the erythroid-restricted uptake profile of native B19V, competes with the virus for entry, and serves as a sensitive probe for receptor expression^11,13^. These studies established that VP1u-dependent uptake is confined to erythropoietin-dependent stages of erythroid differentiation and coincides precisely with the cellular window permissive for productive B19V infection^10,11,14^. Comparative analyses further revealed that VP1u-mediated uptake is conserved among primate erythroparvoviruses, supporting the existence of an evolutionarily conserved cellular receptor^13^. A recent study demonstrated that VP1u-dependent uptake also occurs in villous trophoblasts of the human placenta, showing that expression of a functional VP1uR extends viral entry competence beyond the erythroid lineage and providing a mechanistic basis for how B19V can traverse the placental barrier and reach the fetus^15^.

Despite this extensive functional characterization, the molecular identity of the cellular receptor mediating VP1u-dependent virus uptake has remained elusive. Identifying the VP1uR is essential for understanding the fundamental mechanism that governs B19V tropism and viral entry, both in the target erythroid progenitors and in transplacental transmission. To address this long-standing question, we investigated cell receptors engaged by the VP1u RBD in erythroid cells. Using a combination of proximity labeling, cell-based binding and uptake assays, direct binding experiments, and structural approaches, we have confirmed that human transferrin receptor 1 (TfR1/CD71) is recognized and binds specifically to the VP1u RBD. Furthermore, blocking TfR1 accessibility with the specific antibody OKT9 indicates that B19V interaction with TfR1 is essential for virus uptake.

## Results

### Proximity labeling identifies TfR1 as a VP1u-proximal cell surface protein

To identify host factors in close proximity to VP1u at the cell surface, we applied the enzyme-based Selective Proteomic Proximity Labeling Assay Using Tyramide (SPPLAT)^16^. In this approach, horseradish peroxidase (HRP) generates short-lived tyramide radicals in the presence of H₂O₂, resulting in covalent biotin labeling of proteins within a restricted spatial radius (Extended Data Fig. 1).

Recombinant VP1u was covalently conjugated to HRP, and the integrity of the conjugates was verified biochemically prior to use (Fig. 1A). UT7/Epo cells were incubated with HRP-VP1u at 4 °C to permit surface binding while preventing internalization. Cells exposed to unconjugated HRP and VP1u served as controls. Following activation of the labeling reaction with biotinylated tyramide, biotin deposition was assessed by immunofluorescence. HRP-VP1u and biotin signals showed strong colocalization at the plasma membrane, whereas control conditions yielded only background staining, confirming proximity-dependent labeling at the VP1u binding site (Fig. 1B).

**Figure 1.**
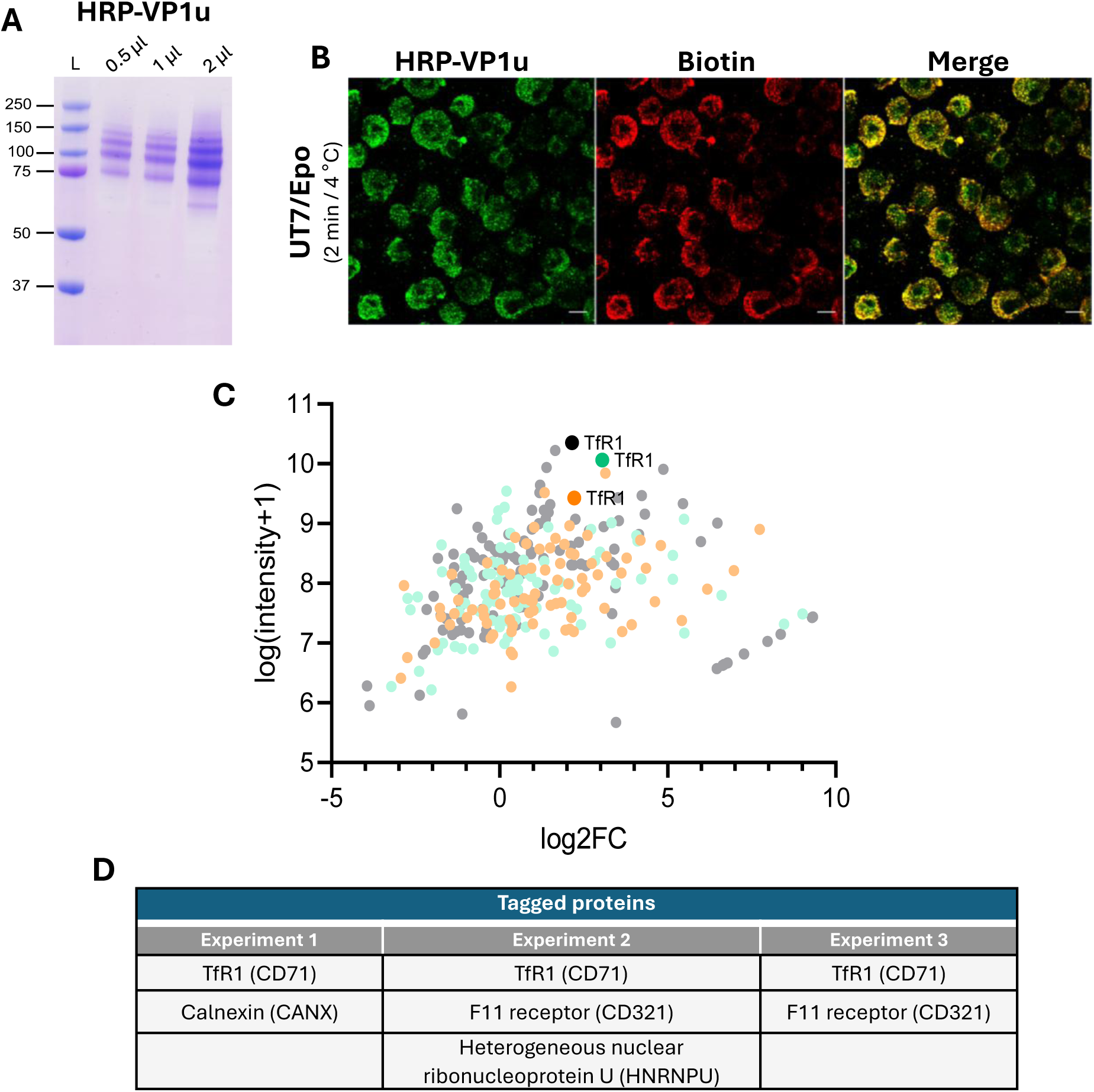
Proximity labeling identifies TfR1 as a VP1u-proximal cell surface protein. **(A)** SDS-PAGE analysis of maleimide-conjugated HRP-VP1u. **(B)** Proximity labeling in UT7/Epo cells. HRP-VP1u was bound to cells at 4 °C, followed by pulse labeling. HRP-VP1u and biotinylated proteins were detected using anti-FLAG and anti-biotin antibodies, respectively. A representative confocal image is shown. Scale bar: 10 μm. **(C)** Mass spectrometry analysis of affinity-purified biotinylated proteins. Proteins identified in three independent experiments were plotted according to log₂ fold change (log2FC) relative to HRP-only controls and protein intensity. Common hits are shown in distinct colors, with TfR1 highlighted. **(D)** Proteins identified as containing tyramide-modified tyrosine residues in the three independent experiments. TfR1 is highlighted in red and represents the only protein detected in all three experiments.

Biotinylated proteins were affinity-purified and subjected to quantitative mass spectrometry. Peptides bearing tyramide-modified tyrosines were prioritized and filtered against the Cell Surface Protein Atlas (CSPA) database to restrict analysis to bona fide plasma membrane proteins^17^. Across three independent experiments, transferrin receptor 1 (TfR1/CD71) was the only reproducibly tyramide-labeled surface protein common to all datasets. Other candidates were either inconsistently detected between replicates or corresponded to proteins without well-established cell surface localization. Comparative analysis of shared proteins based on log₂ fold-change and absolute intensity consistently revealed TfR1 as highly enriched relative to controls (Fig. 1C, D). Together, these data identified TfR1 as the most robust and reproducible VP1u-proximal cell surface protein emerging from the proximity labeling screen.

### VP1u, but not intact B19V, colocalizes with TfR1 at the plasma membrane

To determine whether B19V directly engages TfR1 at the cell surface, we compared the binding pattern of recombinant VP1u with that of intact virions. Recombinant full-length VP1u containing a C-terminal cysteine residue to enable dimerization was expressed and purified as previously described^12^ and detected via its C-terminal FLAG tag. Cells were incubated with recombinant FLAG-tagged VP1u or with native B19V at 4 °C to permit surface binding while preventing endocytosis, thereby restricting the analysis to primary attachment events. TfR1 was visualized using an antibody directed against its cytosolic tail, avoiding potential interference with ligand binding at the ectodomain. Intact virions were detected using a monoclonal antibody specific for conformational epitopes of the capsid.

Confocal microscopy revealed pronounced colocalization of recombinant VP1u with TfR1 at the plasma membrane, as evidenced by strong signal overlap in merged images. In contrast, surface-bound B19V displayed a distinct distribution pattern with no evident colocalization with TfR1 under identical conditions (Fig. 2). These observations indicate that while the VP1u associates with TfR1 at the cell surface, intact virions do not measurably colocalize with the receptor during initial attachment.

**Figure 2.**
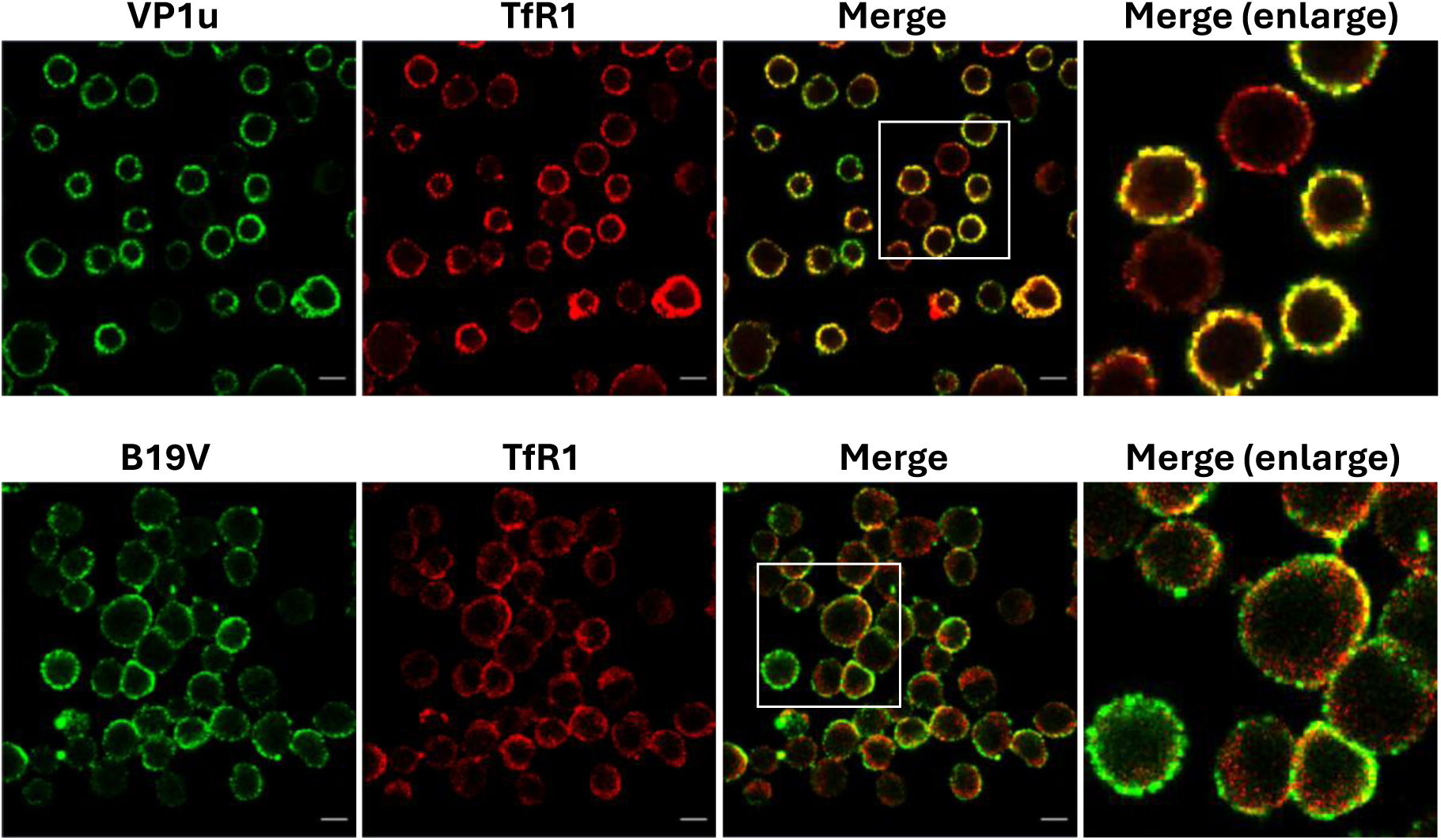
Recombinant VP1u colocalizes with TfR1 at the plasma membrane, whereas intact B19V does not. Confocal microscopy analysis of UT7/Epo cells incubated at 4 °C with recombinant FLAG-tagged VP1u or intact B19V to assess primary surface interactions. Top panels: VP1u (green) and TfR1 (red). Merged images demonstrate substantial colocalization at the cell periphery. Bottom panels: B19V (green) and TfR1 (red). Merged images show distinct spatial distribution with no evident colocalization. Enlarged views highlight the differential overlap patterns. TfR1 was detected using an antibody directed against its cytosolic tail. Recombinant VP1u was detected via its C-terminal FLAG tag, and native B19V was detected using a monoclonal antibody specific for intact capsids (mAb 860-55D). Scale bar: 10 μm.

### Antibody-mediated blockade of TfR1 differentially affects MS2-VP1u and B19V entry

To evaluate the role of TfR1 in VP1u-mediated B19V entry, we used the well-characterized murine monoclonal antibody OKT9, which recognizes the apical domain of human TfR1^18,19,20^. OKT9 has been widely used to inhibit TfR1-dependent viral entry, specifically for viruses that engage the apical domain of TfR1. We therefore used OKT9 to assess whether steric blockade of the TfR1 ectodomain affects VP1u binding and uptake, as well as B19V attachment, internalization, and infection.

Preincubation of UT7/Epo cells with OKT9 completely abolished both binding and uptake of MS2 bacteriophage capsids covalently conjugated to VP1u (MS2-VP1u) via click chemistry, as visualized by confocal microscopy (Fig. 3A). In contrast, in cells treated with the same concentration of an isotype-matched mouse IgG control antibody, MS2-VP1u displayed strong surface binding and internalization. These data demonstrate that OKT9 binding to TfR1 prevents VP1u attachment to and uptake by host cells.

**Figure 3.**
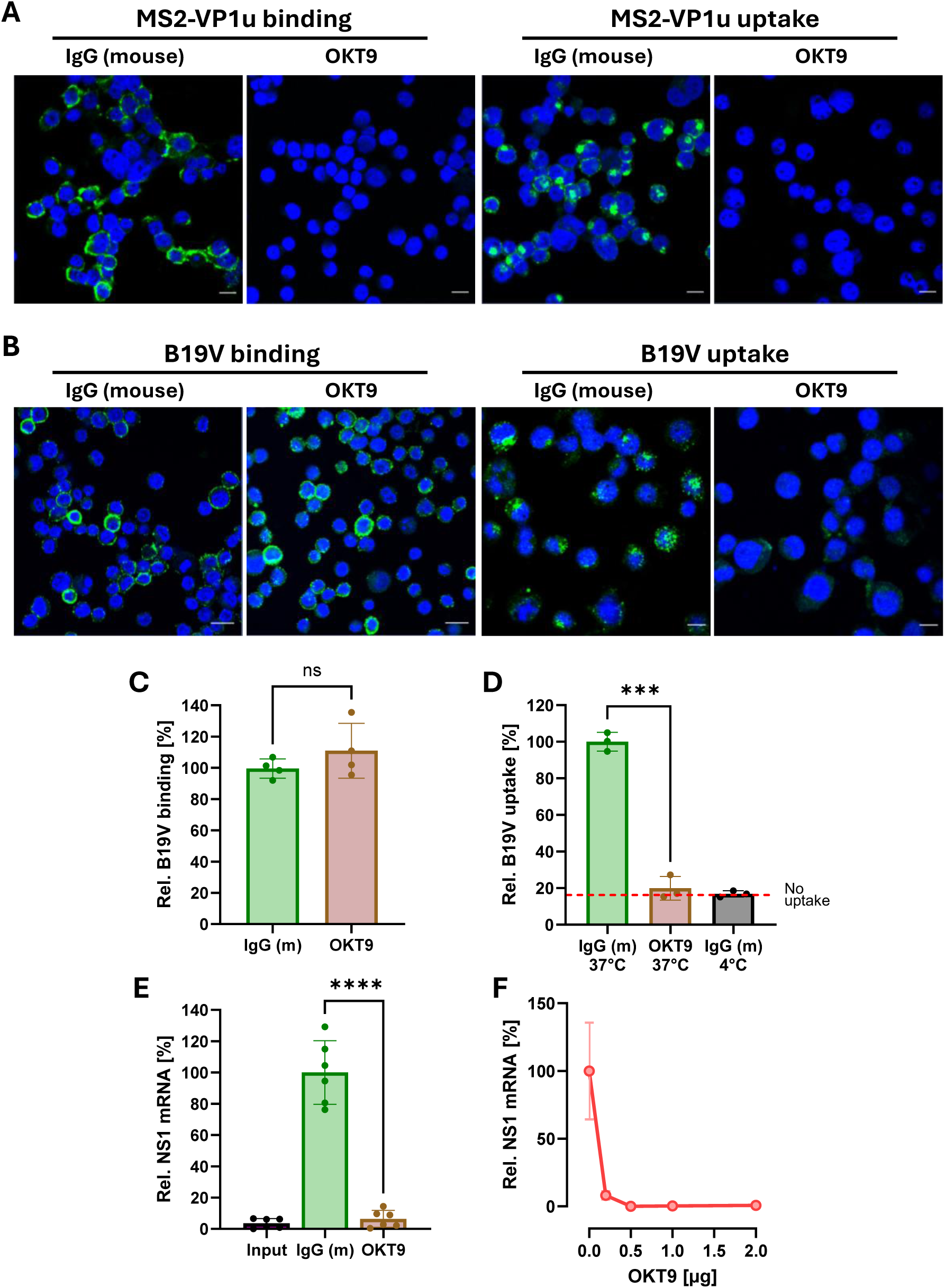
OKT9-mediated TfR1 blockade inhibits VP1u binding, B19V uptake, and infection. **(A)** Confocal microscopy of UT7/Epo cells preincubated with OKT9 or isotype-matched IgG control and exposed to MS2-VP1u-Atto 488 for 30 min at 4 °C (binding) or 37 °C (internalization). Scale bar: 10 μm. **(B)** Confocal microscopy of cells pretreated with OKT9 or IgG and incubated with B19V. B19V was detected using a monoclonal antibody specific to intact capsids (mAb 860-55D). Scale bar: 10 μm. **(C)** Quantification of B19V binding at 4 °C measured by qPCR. **(D)** Quantification of internalized B19V genomes following uptake at 37 °C. Trypsinization was used to remove non-internalized particles. **(E)** NS1 mRNA levels 24 h post-infection following preincubation with 2 µg OKT9. **(F)** Dose-dependent inhibition of NS1 mRNA synthesis following preincubation with increasing concentrations of OKT9 (0, 0.2, 0.5, 1, and 2 µg). All results are presented as the mean ± SD of three or more independent experiments. Statistical significance was calculated using two-sided Student’s t-test. ∗∗∗p < 0.001; ∗∗∗∗p < 0.0001; ns, non-significant.

We next examined intact B19V under analogous conditions. Cells were preincubated with OKT9 for 30 min at 4 °C prior to virus addition. When binding was assessed at 4 °C, OKT9 did not reduce the intensity or distribution of surface-associated virus compared to the IgG control (Fig. 3B), suggesting that initial host cell attachment does not require an interaction with TfR1 apical domains. In contrast, when uptake was permitted at 37 °C for 30 min, OKT9 treatment blocked viral internalization as assessed by confocal microscopy (Fig. 3B). Quantitative PCR analysis of cell-associated and internalized viral genomes corroborated these observations. For this assay, non-internalized particles were removed by trypsinization prior to DNA extraction. The results confirmed that OKT9 blockade does not impair initial binding but prevents subsequent uptake (Fig. 3C, D).

Finally, the requirement for TfR1 interaction in B19V uptake and productive infection was assessed by preincubation of cells with 2 µg OKT9 prior to B19V infection, followed by quantification of NS1 mRNA levels 24 h post-infection. The OKT9 blockade completely inhibited NS1 expression compared with IgG-treated controls (Fig. 3E). Titration of OKT9 (0, 0.2, 0.5, 1, and 2 µg) revealed a dose-dependent inhibition of NS1 mRNA synthesis, reaching near-complete suppression at 0.2 µg (Fig. 3F), indicating that efficient B19V infection depends on TfR1 interaction. Together with the binding and uptake measurements, these data show that TfR1 is not required for initial virion attachment but must be engaged for subsequent internalization. These findings are consistent with a model in which B19V first associates with additional surface factor(s) before VP1u-dependent engagement of TfR1 triggers uptake and productive entry.

### VP1u binding is restricted to erythroid cells despite widespread TfR1 expression

Previous work has consistently demonstrated that VP1u binding and B19V uptake are restricted to erythroid cells, whereas non-erythroid cell types fail to support VP1u binding or viral uptake^14–16^. Following identification of TfR1 as the VP1u-binding receptor, we examined whether this restriction reflects differences in TfR1 surface expression or instead depends on erythroid-specific factors independent of overall receptor abundance.

Non-erythroid cell lines representing distinct tissue origins, including HeLa (epithelial), HepG2 (hepatic), and Jurkat (T lymphoid) cells, were analyzed for TfR1 expression and VP1u binding. In parallel, Ku812Ep6 cells were included as an erythroid-like model. Ku812 cells, derived from a patient with chronic myelogenous leukemia^21^, display erythroid-associated features, can undergo erythroid differentiation in response to erythropoietin^22^. A highly permissive subclone, KU812Ep6, was generated by erythroid enrichment and limiting dilution and has been extensively used, together with UT7/Epo cells, as a model system to study B19V binding, uptake, and infection^23^. Cells were incubated with recombinant FLAG-tagged VP1u at 4 °C to assess surface binding, and TfR1 was detected using an antibody directed against its cytosolic tail following fixation and permeabilization. Confocal microscopy revealed a substantial plasma membrane-associated TfR1 expression in all cell types examined. However, VP1u binding was detected exclusively in KU812Ep6 cells and remained undetectable in non-erythroid cells under identical conditions (Fig. 4A).

**Figure 4.**
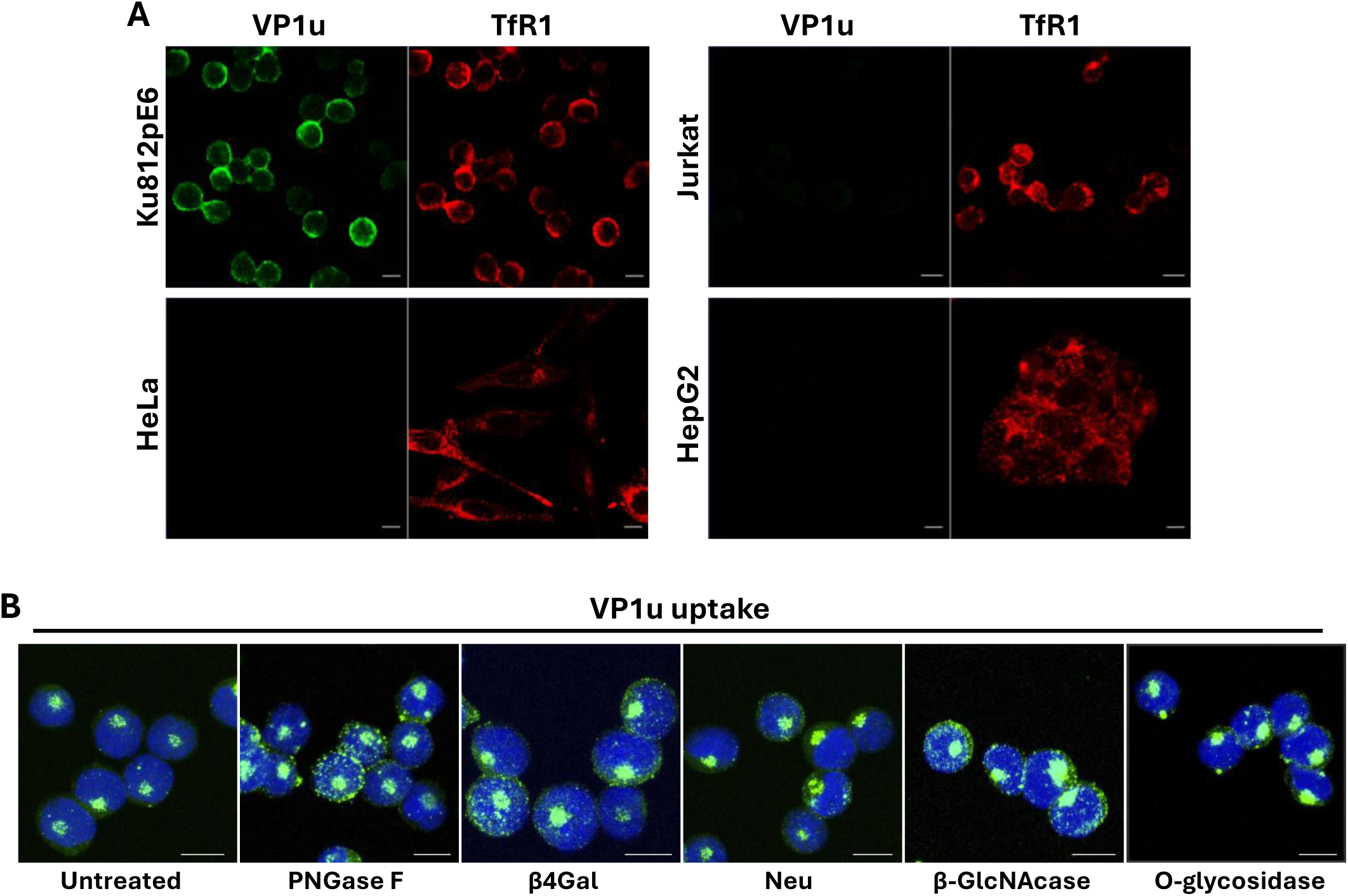
VP1u binding is restricted to erythroid cells and is independent of TfR1 glycosylation. **(A)** Confocal microscopy of erythroid (Ku812Ep6), and non-erythroid cells (HeLa, Jurkat and HepG2 cells) incubated with recombinant FLAG-tagged VP1u at 4 °C. VP1u was detected with anti-FLAG antibody (green) and TfR1 with an antibody against its cytosolic domain (red). **(B)** UT7/Epo cells treated with PNGase F, β1-4-galactosidase (β4Gal), neuraminidase (Neu), β-N-acetylglucosaminidase (β-GlcNAcase), or O-glycosidase prior to incubation with recombinant FLAG-tagged VP1u at 4 °C. VP1u binding was assessed by confocal microscopy. DAPI (blue). Scale bar: 10 μm

These findings indicate that lack of VP1u binding in non-erythroid cells cannot be attributed to absence of TfR1 at the cell surface and instead point to an erythroid-specific context required for productive VP1u-TfR1 interaction.

### TfR1 glycosylation does not affect VP1u uptake in erythroid cells

Because TfR1 is broadly expressed across diverse cell types whereas B19V selectively infects erythroid cells, we considered whether differences in receptor glycosylation might influence VP1u recognition. UT7/Epo cells were treated with glycosidases targeting defined carbohydrate moieties prior to incubation with recombinant FLAG-tagged VP1u. PNGase F was used to remove N-linked glycans, whereas β1-4-galactosidase, neuraminidase, and N-acetylglucosaminidase were used to trim terminal galactose, sialic acid, and N-acetylglucosamine residues, respectively. Confocal microscopy analysis showed that enzymatic removal of these carbohydrate structures did not reduce VP1u uptake (Fig. 4B).

Although the efficiency of enzymatic trimming was not independently quantified, the absence of any detectable reduction in VP1u uptake following treatment with multiple glycosidases indicates that accessible surface glycans are not essential determinants of the VP1u-TfR1 interaction in erythroid cells. These results therefore argue against differential TfR1 glycosylation as the basis for the erythroid restriction of VP1u binding and instead point to additional cell type-specific determinants beyond TfR1 glycan composition.

### VP1u binds to TfR1 in a cell-free system

To assess the VP1u-TfR1 interaction in a cell-free condition and determine the binding affinity between RBD and the receptor, we performed mass photometry and bio-layer interferometry (BLI) analysis. Monomeric and dimeric VP1u proteins were incubated with TfR1 (final concentration 3 µM) at a ratio of 2:1 and 1:1, respectively, for mass photometry experiment, which confirmed the formation of the VP1u-TfR1 complex (Extended Data Fig. 2).

For the BLI analysis, Fc-tagged TfR1 was immobilized on Fc-capture biosensors, and the association and dissociation kinetics of VP1u monomer and dimer were measured using serial dilutions (Extended Data Fig. 3). The resulting graphs showed concentration-dependent binding kinetics; however, the data were not consistent with a 1:1 Langmuir binding model for both samples. Instead, the data were best fitted using a heterogeneous ligand (2:1) model, which yielded R^2^ values higher than 0.98, suggesting the presence of two different binding populations on the sensor surface. The calculated K_D_1 (K_D_2) values were about 3.1 (3.7) µM for the TfR-VP1u monomer and 2.6 (4.5) µM for the TfR1-dimer, respectively (Extended Data Fig. 3). We did not observe significant avidity effects in the TfR1-VP1u dimer interaction, suggesting the VP1u dimer did not engage dimeric TfR1 in a bivalent manner and did not cross-link two TfRs molecules.

Consistent with these observations, experiments in UT7/Epo cells showed only a modest increase in surface binding of the VP1u dimer compared with the monomer when binding was measured at 4 °C. In contrast, assays performed at 37 °C, which allow receptor-mediated uptake, revealed a pronounced advantage of the VP1u dimer (Extended Data Fig. 4). These findings are in agreement with previous work demonstrating that VP1u dimers bind and internalize more efficiently than monomeric VP1u^12^.

Together, these results indicate that dimerization does not substantially increase the intrinsic affinity of VP1u for TfR1 but enhances functional interaction with the receptor in a cellular context. One possible explanation is that VP1u dimerization facilitates more efficient engagement of TfR1 at the cell surface, potentially involving local receptor clustering or cooperative interactions that can occur in the membrane environment. Such effects would not be captured in the BLI assay, where accessible TfR1 molecules are immobilized on the biosensor surface and lack lateral mobility.

### Cryo-EM 3D reconstruction reveals VP1u binding to TfR1 apical domain

To elucidate the molecular interaction between VP1u and TfR1 and determine the three-dimensional structure of the complex, we performed cryo-EM single-particle analysis.

Because monomeric VP1u interacts with dimeric TfR1, it was likely that two VP1u monomers would bind to each TfR1 dimer. Monomeric VP1u was therefore used to avoid cross-linking of TfR1 through bivalent binding by VP1u dimers. Full-length VP1u was incubated with the recombinant human TfR1 ectodomain at a 2:1 molar ratio, and the resulting complex was vitrified for cryo-EM data collection and single-particle reconstruction (Extended Data Table 1).

The three-dimensional map of the VP1u-TfR1 complex was initially reconstructed at 2.23 Å resolution using 681,957 particles with C2 symmetry (Extended Data Fig. 5). In parallel, the human TfR1 ectodomain alone was vitrified and analyzed, yielding a structure at 2.42 Å resolution (Extended Data Fig. 6). Both maps revealed the canonical TfR1 architecture, characterized by a butterfly-shaped dimeric ectodomain composed of apical, protease-like, and helical domains. Notably, in the VP1u-TfR1 complex map, two additional unfilled densities were observed, one on the side of each apical domain, which were of dimensions that could correspond to the receptor-binding domains (RBDs) of VP1u (Extended Data Fig. 5 and Fig. 5A).

**Figure 5.**
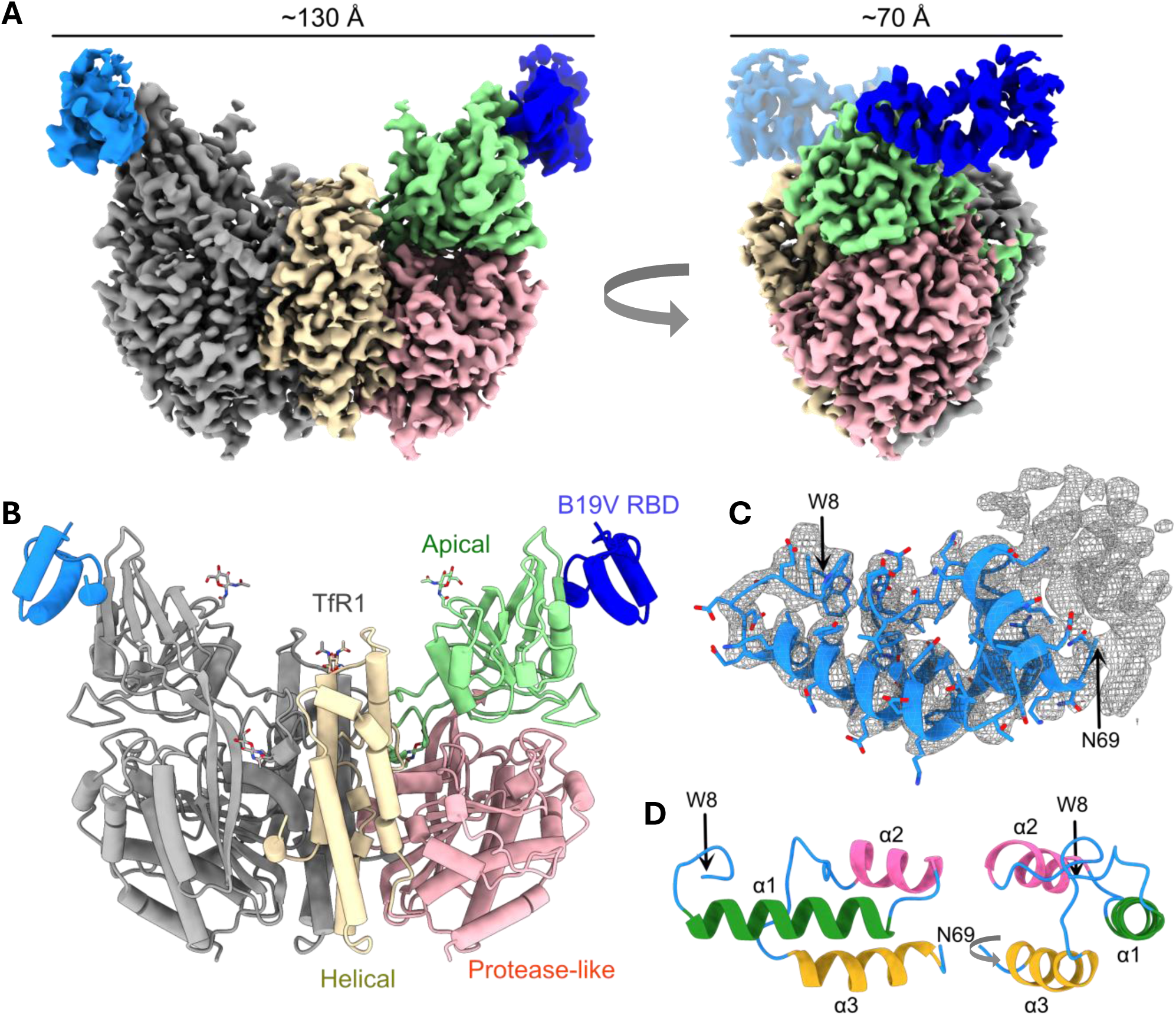
VP1u RBD binds to the apical domain of TfR1. **(A)** Cryo-EM 3D reconstruction of TfR1 in complex with recombinant VP1u monomer. One TfR1 monomer is colored to distinguish the apical, protease-like, and helical domains. The two additional densities corresponding to the two bound VP1u molecules are shown in blue and light blue. **(B)** Atomic model of the TfR1-VP1u RBD complex structure, depicted as a ribbon diagram with cylinders and stubs representing α-helices and β-strands, respectively. **(C)** *De novo* model of RBD (residues 8-69) with cryo-EM density, shown as a mesh envelope. **(D)** The atomic model of the VP1u RBD displayed as a ribbon diagram. The three α-helices are highlighted and labeled.

Although the TfR1 density was well defined, the additional densities exhibited weaker signal intensity and lower local resolution, particularly in their distal regions, suggesting partial occupancy or structural flexibility. To improve the putative RBD density, particles were symmetry-expanded and subjected to two rounds of focused classification for each RBD. The resulting reconstruction showed enhanced RBD density, albeit at a slightly lower overall resolution of 2.47 Å. Furthermore, the classified symmetry-expanded particles were used for three-dimensional flexible reconstruction, which substantially improved the overall quality and continuity of the density corresponding to bound VP1u-RBD (Extended Data Fig. 5 and Fig. 5A).

### Atomic model building identifies VP1u RBD

We refined the crystal structure of TfR1 against our TfR cryo-EM density map to obtain the final model (Extended Data Fig. 7A)^24^. Comparison of the two atomic models revealed conformational differences in the apical traverse loop, which spans the interface between the apical and protease-like domains, and in C-terminal tail of the polypeptide chain, with local Cα RMSDs of 5.543 Å and 4.630 Å, respectively (overall Cα RMSD: 1.669 Å)(Extended Data Fig. 7B-D). Both regions coordinate Sm^3+^ ions in the crystal structure^24^, likely reflecting the crystallization experimental conditions.

The TfR1 model obtained from the cryo-EM map was then refined against the VP1u-TfR1 complex density map (Fig. 5B). The refined TfR in complex with VP1u revealed local conformational changes in the apical loop that engages VP1u when compared with the TfR-alone structure (overall Cα RMSD: 0.664 Å)(Extended Data Fig. 7E and F). After finalizing the TfR1 model, the remaining unassigned density was used for *de novo* model building with ModelAngelo^25^, guided by the amino acid sequences of the VP1u RBD (residues 1-71). We also tested the full-length VP1u sequence (residues 1-227). Both sequences generated models with identical amino acid registration, which were further refined to produce an atomic model corresponding to residues 8-69 that was built into the density with high confidence (Fig. 5C). Consistent with previous predictions^12^, the VP1u receptor-binding domain contains three α-helices (α1– α3) that assemble into a compact three-helix bundle fold (Fig. 5D). No density was observed N-terminal to Trp8, whereas weak and disordered density extended from Asn69 (Fig. 5C). The remaining C-terminal region of VP1u, including the phospholipase A2 (PLA_2_) domain, was not resolved in the map. It is possible the flexible C-terminal region influenced the binding of another VP1u on the opposite side of the TfR, which may explain the non-Langmuir binding behavior we observed in BLI analysis (Extended Data Fig. 3).

Three glycosylation sites exhibited clear density corresponding to the first glycan residues at Asn251, Asn317, and Asn727 (Fig 5B), although more extended glycan density was observed only at lower, noisier contour levels (Extended Data Fig. 5 and 6). None of these glycosylation sites were positioned close enough to interact with the VP1u receptor-binding domain. We also analyzed the VP1u-TfR1 complex structure using PRODIGY web server^26^ to predict the binding affinity between RBD and TfR1, yielding a K_D_ of 1 μM, which is within the same order of magnitude as the BLI measurements. Overall, the cryo-EM structure of the VP1u-TfR1 complex demonstrated that two VP1u RBDs bind, one to each TfR1 apical domain on each side of one TfR1 molecule.

**Figure 6.**
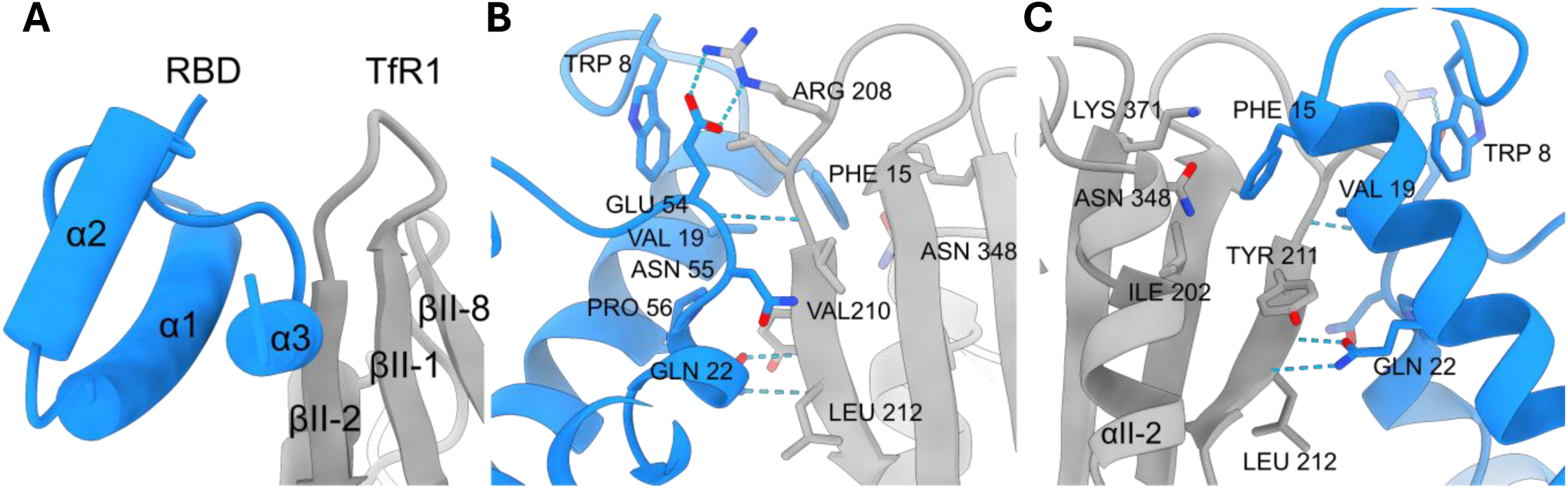
Molecular interaction between VP1u RBD and TfR1. **(A)** Close-up view of the RBD-TfR1 binding interface from Fig. 5C. Secondary structure elements of the RBD and TfR1 are labeled. **(B and C)** Contacting residues between RBD and TfR1 are shown as stick representations with side chains displayed and residues labeled.

### Molecular interactions between RBD and TfR1

Using the atomic model of the VP1u-TfR1 complex, we identified the contact residues between the RBD and the receptor. The N-terminal loop, BC loop, and the A and C α-helices of the RBD interacted with the apical domain of TfR1(Fig. 6A), primarily through the βII-1-βII-2 loop and the βII-2 strand (residues 208-212)(Fig. 6B), as well as three residues from the βII-1 strand, βII-8 strand, and αII-2 helix (I202, K371, and N348, respectively)(Fig. 6C). Notably, Phe15 of the RBD is the only residue that contacts all three TfR residues outside the βII-2 strand, whereas the remaining RBD residues interact exclusively with the βII-1-βII-2 loop and the βII-2 strand of TfR (Fig. 6C).

Previously through mutational analysis, we identified a cluster of amino acids that play critical roles in VP1u internalization^12^. Mapping these functionally important residues onto the RBD-TfR1 structure revealed that many residues, including WW8/9, F15, V19, Q22, and EN54/55, are located at the receptor-binding interface, whereas the remaining residues are positioned at the helical interface, where mutations are likely to affect RBD stability (Fig 7A). Notably, most of the contact residues identified in the RBD-TfR1 complex structure, except for one untested residue (Pro 56)(Fig 6B), exhibited significantly reduced VP1u activity when mutated^12^, supporting the conclusion that the previously unidentified VP1u receptor (VP1uR) in our earlier study was in fact, TfR1.

**Figure 7.**
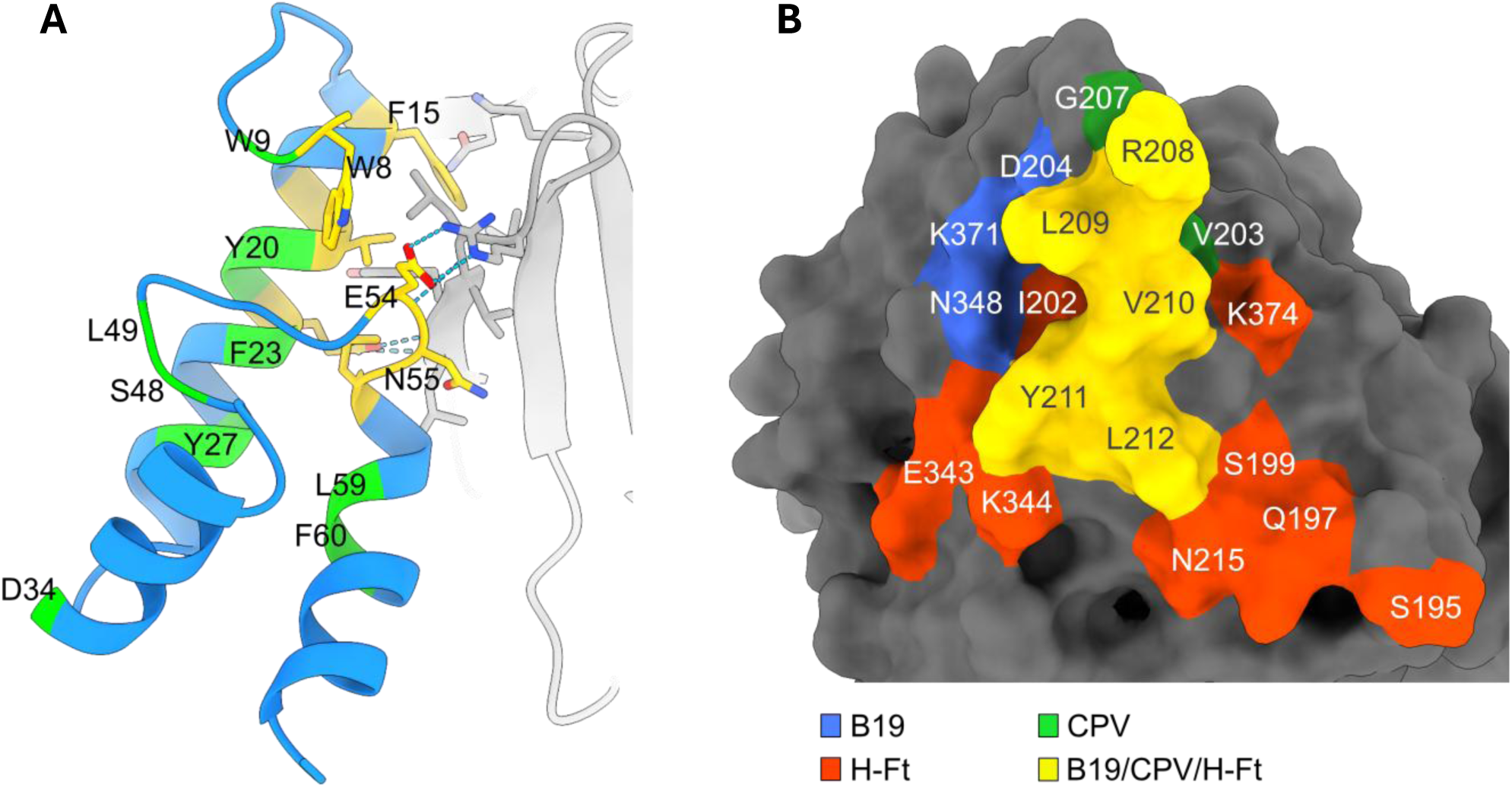
Mapping of critical residues. **(A)** Residues previously examined for their functional roles in VP1u internalization are highlighted. Residues located within the contact interface with TfR1 are colored yellow, whereas those not found at the interface are colored green. **(B)** Binding sites of the B19V RBD, human ferritin, and canine parvovirus (CPV) are mapped on the surface of human TfR1. Each interaction site is colored according to the key shown at the bottom. Residue numbering of canine TfR was aligned with that of human TfR. Only consistent contacts for CPV is shown^27^.

We then compared the RBD contact residues with those involved in interactions with human ferritin and canine parvovirus (Fig. 7B). All three molecules commonly engaged residues 208-212 on the βII-1-βII-2 loop and the βII-2 strand. The human ferritin and CPV binding sites involve broader surface areas, likely reflecting the larger, spherical shapes of these molecules, although CPV-TfR1 contacts are transient^27^. In contrast, the interaction interface between the VP1u RBD and TfR1 covers a smaller area about 480 Å². The residues on the βII-1 and βII-8 strands and the αII-2 helix (I202, K371, and N348) are unique to B19V contacts. Residue K371 is also found among the contact residues of other human pathogens, including Machupo virus (MACV)^28^ and the parasite Plasmodium vivax^29^.

## Discussion

B19V is characterized by a remarkably strict tropism for EPCs, a restriction that underlies its pathogenesis^30^. Extensive work over the past decade established that this narrow tropism is primarily controlled at the level of viral uptake and is mediated by VP1u, specifically via an RBD located within its N-terminus ^10,13,14^. VP1u RBD is necessary and sufficient to drive erythroid-restricted internalization, and its uptake profile precisely matches the differentiation window permissive for productive infection^11^. Despite this detailed functional characterization, the molecular identity of the VP1u receptor has remained unresolved. AXL was recently proposed to contribute to B19V entry into EPCs. However, AXL expression is not restricted to erythroid cells, and inhibition or ectopic expression experiments indicate that AXL alone is not sufficient to confer full susceptibility to B19V infection. These observations suggest that AXL may act as a contributory entry factor rather than the primary receptor responsible for VP1u-mediated uptake^31^.

Here, we identify TfR1 (CD71) as a specific binding partner of VP1u and show that it corresponds to the previously postulated VP1u receptor (VP1uR) required for VP1u-mediated viral internalization. Proximity labeling reproducibly identified TfR1 as the only consistently enriched cell surface protein across independent experiments. Recombinant VP1u colocalized with TfR1 at the plasma membrane under conditions restricting analysis to primary binding events, whereas intact virions did not show detectable colocalization at this stage. Functional blockade of the TfR1 apical domain with OKT9 abolished VP1u binding and uptake and prevented B19V internalization and infection, while virus attachment at 4 °C was not affected. The inhibitory effect was dose-dependent and nearly complete at low antibody concentrations, further supporting the functional relevance of this interaction. In parallel, cell-free binding assays confirmed the VP1u-TfR1 binding and measured the binding kinetics. Cryo-EM analysis demonstrates direct binding of the VP1u RBD to the TfR1 ectodomain and defines the molecular interface at near-atomic resolution. Residues previously shown by mutagenesis to be critical for VP1u uptake localize directly to the receptor-binding surface in the structure, providing independent validation of the identified interface^12^. The convergence of proteomic enrichment, antibody blockade, uptake assays, infection readouts, biochemical assays, and structural characterization provides orthogonal evidence that TfR1 is the receptor engaged by VP1u to trigger B19V internalization.

The differential effect of TfR1 blockade on attachment versus internalization supports a multistep mechanism for B19V entry. The observation that virion attachment occurs independently of TfR1, whereas internalization requires TfR1 engagement, indicates that intact B19V first associates with the erythroid cell surface through VP1u-independent interaction(s) before engaging TfR1. This sequential mechanism is consistent with our previous demonstration that VP1u is not exposed on the native capsid surface but becomes accessible only after contact with permissive erythroid cells^32^. More recently, we showed that patient-derived B19V particles circulate coated with host-derived protease inhibitors that dissociate upon interaction with erythroid cells, accompanied by structural rearrangements and exposure of the otherwise internal VP1u region^33^. These findings support a priming-dependent entry mechanism in which native virions initially attach to the host through capsid-exposed determinant(s) independent of TfR1, followed by conformational remodeling that enables VP1u engagement of TfR1, which subsequently triggers internalization. The identity of the primary attachment factor remains to be defined, but the present data separate the attachment step from the TfR1-dependent uptake step.

A second central question concerns the restriction of VP1u-TfR1 interaction to erythroid cells despite widespread TfR1 expression. The present data impose important constraints. In previous studies and in this study, multiple non-erythroid cell lines express TfR1 yet do not support VP1u binding^10,13^. Enzymatic removal of N-linked glycans and terminal carbohydrate residues from UT7/Epo cells does not impair VP1u binding. Furthermore, the cryo-EM structure demonstrates direct binding of VP1u to the recombinant TfR1 ectodomain produced in HEK293 cells, indicating that non-erythroid-derived TfR1 is intrinsically capable of interaction. This apparent discrepancy indicates that the inability of VP1u to bind non-erythroid cells does not reflect an intrinsic incompatibility of the receptor but instead arises from the cellular context in which TfR1 is presented at the cell surface. Together, these observations argue against models based on erythroid-specific TfR1 isoforms or glycosylation-dependent recognition.

Structural mapping reveals that B19V targets a defined hotspot within the apical domain of TfR1. This region is also engaged by unrelated pathogens, including New World arenaviruses^20^ and carnivore parvoviruses^27,34,35^, underscoring the apical surface as a recurrent entry interface. In contrast to carnivore parvoviruses, B19V mediates receptor binding through a specialized capsid extension (VP1u) rather than the icosahedral capsid topology^27^. The VP1u-TfR1 structure therefore illustrates how a modular capsid element can govern cell-type-restricted uptake while the structural shell remains functionally distinct.

In summary, we have established the VP1u specific recognition and binding of TfR1. Furthermore, the combined proteomic, functional, and structural evidence indicates TfR1 is the previously unknown VP1u receptor that is required for VP1u-dependent internalization of B19V. These data resolve a long-standing question in B19V biology by identifying the molecular receptor that mediates erythroid-restricted uptake and by defining its structural interface. The clear separation between primary attachment and TfR1-dependent internalization reveals the sequential organization of viral entry. Together, these findings provide a mechanistic framework for B19V uptake and establish TfR1 as the entry receptor that enables internalization into target progenitor cells.

## Materials and Methods

### Cells and viruses

UT7/Epo cells, provided by E. Morita (Tohoku University School of Medicine, Japan) were cultured in Eagle’s Minimum Essential Medium (MEM, Thermo Fisher Scientific, Waltham, MA, USA) with 5% FCS and 2 U/mL Epo. KU812Ep6 and Jurkat cells (ATCC; TIB-152) were maintained in RPMI 1640 with 10% FCS; KU812Ep6 cultures were additionally supplemented with 6 U/mL Epo. The hepatocarcinoma cell line HepG2 (ATCC; HB-8065) and HeLa cells (ATCC; CCL-2) were cultured in DMEM with 10% FCS. All cells were routinely maintained at 37 °C and 5% CO_2_.

Native B19V was obtained from de-identified plasma samples of infected, seronegative individuals, confirmed by virus-specific serology (CSL Behring, Bern, Switzerland). Infected plasma was thawed and clarified by centrifugation at 4,000 rpm for 10 minutes at 4 °C. To generate IgG-depleted B19V, the plasma was subjected to 20% (w/v) sucrose/PBS ultracentrifugation using a TLA100.3 rotor (Beckmann) at 195,000 × g at 4 °C for 2 hours. The viral pellet was resuspended in PBS and contaminant human IgG was removed by incubation with Protein G Plus Agarose (sc-2002, Santa Cruz Biotechnology) at 4 °C for 1 hour. Viral genomes were quantified by quantitative PCR (qPCR) using Luna Universal One-Step Reaction Mix (M3003, New England Biolabs, NEB, Ipswich, MA, USA) with primers specific to the NS1-coding region of the genome as mentioned below. Plasmids containing the complete B19V genome were used as external standards in 10-fold serial dilutions.

### Recombinant VP1u constructs and MS2-VP1u VLPs

Recombinant full-length VP1u was expressed and purified as previously described^12,14^. Briefly, *E. coli* BL21(DE3) cells were transformed with pT7-FLAG-MAT-Tag-2 plasmids encoding either full-length VP1u or a modified VP1u containing an additional C-terminal cysteine residue to allow disulfide-mediated dimerization. The resulting proteins therefore yielded either monomeric VP1u or dimerizable VP1u variants, enabling comparison of monomeric and dimeric forms in subsequent experiments. Expression and purification of recombinant VP1u proteins were performed as previously described^14^.

For proximity labeling studies, a pT7-FLAG-MAT-Tag-2 vector encoding a C-terminally truncated VP1u of B19V (amino acids 1-99) was transformed into *E. coli* BL21(DE3) cells. This construct lacks the C-terminal 128 amino acids of the full-length VP1u (227 aa) while retaining a fully functional RBD^12^. A C-terminal cysteine residue was introduced to allow disulfide-mediated dimerization of the protein. Expression and purification of truncated VP1u proteins were performed as previously described^14^. Recombinant VP1u was conjugated to horseradish peroxidase (HRP). The recombinant VP1u was reduced by addition of 5 mM TCEP (Lucerna-Chem, P1021) for 30 min followed by addition of HRP-Maleimide (ImmunoChemistry Technologies, 6294), keeping the VP1u at a 30-fold molar excess. After 16 h at 4 °C, the reactants were separated using an ÄKTA size exclusion column (Superdex 200 increase 10/300) and fractions of 400 µl were collected. Collected proteins were separated by SDS-PAGE and the fractions containing the desired construct were pooled and subsequently concentrated using spin columns with a 30 kDa molecular weight cut-off (Merck Millipore, UFC503008). Concentrated HRP-VP1u was stored at -80 °C.

MS2-bacteriophage virus-like particles (VLPs) were expressed using *E. coli* BL21(DE3), and purified by ultracentrifugation through a 20% sucrose cushion as described elsewhere^13^. MS2-VLPs were purified and labeled with NHS-Atto 488 (Atto-Tec, Siegen, Germany) using a 40-fold molar excess of dye. The reaction was quenched, and labelled VLPs were recovered by centrifugation through a 20% sucrose cushion to remove unreacted dye. Subsequently, MS2 VLPs were incubated with 500-fold molar excess of maleimide-PEG_24_-NHS (22114, Thermo Fisher Scientific) for 1 h. Excess crosslinker was eliminated using a 40 kDa MWCO desalting column. The activated VLPs were then conjugated to reduced recombinant full-length VP1u proteins and further purified by a second 20% sucrose cushion.

### Proximity-based biotinylation of VP1u-proximal proteins

All the steps were performed at 4 °C unless stated otherwise. For each experiment 1.5 *×* 10^7^ cells were washed twice with PBS at room temperature and then resuspended in PBS to reach a final concentration of 5 *×* 10^6^ cells per ml. HRP-VP1u (5 µg/ml) was added and allowed to interact with the cells for 2 h under constant agitation. In parallel, a control sample was incubated with 4 µg/ml VP1u and 1 µg/ml HRP (unconjugated). Cells were pelleted at 700 *×* g for 5 min, washed once with ice-cold PBSA 0.2 % (PBS containing 0.2 % albumin), then resuspended in freshly prepared labelling buffer (50 mM Tris-HCl pH 7.4, 0.03 % H_2_O_2_, 80 µg/ml Tyramide-Biotin [Iris Biotech, LS-3570.0250, 10 mg/ml in DMSO]) and incubated for 2 min. Catalase (Sigma-Aldrich, C9322) was added to a final concentration of 100 U/ml to stop the reaction. Cells were then washed three times with ice-cold PBSA 0.2 %, followed by lysis using ice-cold lysis buffer (20 mM Tris-HCl pH 7.4, 5 mM EDTA pH 8.0, 150 mM NaCl, 1 % Triton X-100, 0.1 M sodium thiocyanate, 1× Protease inhibitor cocktail (500 µl per 5 *×* 10^6^ cells; 11836153001, Roche Diagnostics GmbH, Mannheim, Germany). Samples were vortexed briefly and incubated for 15 min before addition of Benzonase (Millipore, E1014) and incubation for 30 min. Samples were spun at 12’000 *×* g for 10 min to remove insoluble material. In the meantime, neutravidin beads (Thermo Fisher Scientific, 29200) were washed once with PBSA 1 % and twice with lysis buffer. The lysate supernatant was added to the beads and incubated for 1 h under constant agitation. The beads were then washed four times with wash buffer 1 (10 mM Tris-HCl pH 7.4, 1 % Triton X-100, 1 mM EDTA pH 8.0, 0.5 % SDS, 500 mM NaCl, 0.1 M sodium thiocyanate, 1× Protease inhibitor cocktail) and twice with wash buffer 2 (10 mM Tris-HCl pH 7.4, 1 % Triton X-100, 1 mM EDTA pH 8.0, 0.5 % SDS, 0.1 M sodium thiocyanate, 1× Protease inhibitor cocktail). Beads were subsequently resuspended in 4X LDS sample buffer (Thermo Fisher, Waltham, MA).

### Identification of VP1u-proximal proteins

Purified biotinylated proteins were separated on an SDS-PAGE. After the gel front had migrated roughly 1.5 cm, the run was stopped, and proteins were stained using Coomassie gel stain (Thermo Fisher Scientific, 24615). A sterile scalpel was used to cut the resolved proteins of each lane into three horizontal slices. The slices were collected and stored in separate 1.5 ml tubes. Gel pieces were overlaid with 80 % ethanol and stored at 4 °C. Proteins were identified by MS analysis at the Core Facility Proteomics & Mass Spectrometry in Bern. Quantification was performed using the sum of the three most intense peptide intensities (Top3). Variance Stabilizing Normalization (VSN) was employed before summing up of the peptides for Top3. Enrichment of protein was calculated as log_2_ fold-change (log2FC) compared to the control. Additionally, peptides with tyramide-modified tyrosines were identified.

### Binding and internalization assays

Cell surface binding of recombinant VP1u (100 ng), MS2-VP1u particles and B19V (10⁵/cell) was carried out by incubating cells in PBS (pH 7.2) at 4 °C for 30 min. After incubation, cells were washed three times with ice-cold PBS and either subjected to DNA isolation followed by qPCR analysis or fixed for immunofluorescence microscopy. For internalization assays, MS2-VLPs, or B19V were incubated at 37 °C for 30 min. Cells were then washed with PBS. For qPCR-based analysis, cells were subsequently treated with trypsin/EDTA at 37 °C for 4 min to remove surface-associated particles, washed again, and processed for DNA extraction. For immunofluorescence analysis, cells were washed after internalization and processed directly for staining without trypsin treatment. Genomic DNA was purified using the GenCatch Plus Genomic DNA Miniprep Kit (1660250, Epoch Life Science, Missouri City, TX, USA) following the manufacturer’s protocol. Quantitative PCR was performed using the primers B19-F 5′-GGGGCAGCATGTGTTAAG-3′ and B19-R 5′-CCATGCCATATACTGGAACAC-3′.

### Antibody-mediated blockade of the VP1u-TfR1 interaction using OKT9

UT7/Epo cells were preincubated with 2 µg, unless otherwise indicated, of anti-human TfR1 monoclonal antibody OKT9 (103101-0N; Caprico Biotechnologies, Duluth, MN) or with an isotype-matched mouse IgG control (14-4714-82; Thermo Fisher) for 30 min at 4 °C prior to addition of MS2-VP1u or B19V. For binding analyses, antibody-treated cells were incubated with MS2-VP1u or B19V at 4 °C for 30 min, washed extensively with cold PBS, and either fixed for confocal microscopy or processed for DNA extraction and qPCR. For uptake experiments cells were incubated at 37 °C for 30 min to allow internalization. For samples analyzed by qPCR, surface-bound particles were removed by trypsin/EDTA treatment (4 min, 37 °C), followed by washing prior to DNA extraction. For samples analyzed by immunofluorescence, cells were washed after internalization and fixed directly without trypsin treatment. To assess productive infection, UT7/Epo cells were preincubated with OKT9 (0-2 µg) before infection with B19V. Total RNA was isolated 24 h post-infection, and NS1 mRNA levels were determined by qRT-PCR using the following primers: NS1-F: 5′-GGGGCAGCATGTGTTAAG-3′, B19-NS1-R: 5′- CCATGCCATATACTGGAACAC-3′.

### Immunofluorescence microscopy

Colocalization of VP1u with TfR1 was assessed in UT7/Epo cells by incubating recombinant B19 VP1u at 4 °C for 30 min in the presence of a monoclonal rabbit anti-FLAG IgG antibody (14793S, Cell Signalling Technologies, Danvers, MA, USA). The cells were washed several times with ice-cold PBS before fixation in a 1:1 mixture of methanol:acetone at -20 °C for 4 min. The FLAG antibody was detected with a polyclonal Alexa Fluor® 488 conjugated goat anti-rabbit IgG (Thermo Fisher). TfR1 was detected with a mouse anti-human TfR1 (13-6800; Thermo Fisher) specific to residues 3-28 of the human TfR1 cytoplasmic tail and a secondary polyclonal goat anti-mouse IgG conjugated to Alexa Fluor® 594 (Invitrogen). B19V capsids were detected using a monoclonal human IgG anti-B19 VP2 antibody (860-55D, Mikrogen, Neuried, Germany) and a secondary polyclonal goat anti-human IgG conjugated to Alexa Fluor® 488 (Invitrogen). All antibodies were diluted in 2% milk/PBS and carried out at 4 °C, after which specimens were treated with 2 mM CuSO_4_/50 mM NH_4_. The samples were mounted with EMS shield mount (17985-09, Electron Microscopy Sciences, Hatfield, PA, USA) supplemented with 0.2 ng/mL DAPI when indicated. The samples were visualized with a 63× oil immersion objective by laser scanning confocal microscopy (LSM880, Zeiss, Oberkochen, Germany).

### Enzymatic removal of cell surface glycans and VP1u binding assay

To evaluate whether glycosylation of TfR1 contributes to VP1u binding, UT7/Epo cells were subjected to enzymatic removal of specific surface glycans prior to incubation with recombinant VP1u. A total of 3 × 10⁵ cells per condition were washed once with PBS and resuspended in 30 μl of the corresponding reaction buffer. Cells were incubated for 1 h at 37 °C with one of the following enzymes under buffer-matched conditions: PNGase F (500 U; 50 mM sodium phosphate, pH 7.5; P0704, NEB), O-glycosidase (10 U; 50 mM sodium phosphate, pH 7.5; P0733, NEB), β1-4-galactosidase (10 U; 50 mM sodium acetate, pH 5.5, 5 mM CaCl₂; P0745, NEB), β-N-acetylglucosaminidase (10 U; 50 mM sodium acetate, pH 5.5, 5 mM CaCl₂; P0744, NEB), or α2-3,6,8 neuraminidase (40 U; 50 mM sodium acetate, pH 5.5; P0720, NEB). For removal of O-linked glycans, cells were first treated with neuraminidase, washed, and subsequently incubated with O-glycosidase (10 U; 50 mM sodium phosphate, pH 7.5; P0733, NEB). Following enzymatic treatment, cells were washed with PBS and incubated with recombinant FLAG-tagged VP1u at 37 °C for 1 h and processed for immunofluorescence analysis as described above.

### Mass photometry and Bio-layer interferometry

The purified VP1u proteins were incubated with tag-free TfR1 (Acro Biosystems) at room temperature for 30 min, at a final concentration of 4 µM TfR1 and 8 µM VP1u monomer or 4 µM VP1u dimer. The mass of the incubations was measured by using TwoMP (Refeyn). Molecular weights were calibrated using MassFerence P1 calibrant (Refeyn).

For BLI analysis, anti-hIgG Fc capture (AHC) biosensors (Sartorius) were used to bind Fc-tagged TfR1 (Acro Biosystems). Binding kinetics and affinity were measured using Octet R2 (Sartorius). The AHC biosensors were first blocked and hydrated in kinetics buffer (PBS with 0.01 % ovalbumin and 0.02 % Tween 20), which was used during the whole experiments. Binding experiments were performed with the following protocol: 60 s equilibration, 300 s of loading 10 μg/mL of TfR1, 180 s of baseline, 300 s of association with VP1u molecules, and 300 s of dissociation. The data were analyzed using Octet Analysis Studio with a 2:1 (heterogeneous ligand) binding model.

### Sample preparation and Cryo-EM data collection

The purified VP1u proteins were incubated with tag-free TfR1 (Acro Biosystems) at room temperature for 30 min, at a final concentration of 7 µM. Aliquots of 3.5 µl of the incubated sample and TfR1 alone were applied to freshly glow discharged Quantifoil R 1.2/1.3 300 mesh copper grids. The grids were plunged into liquid ethane maintained at liquid nitrogen temperature, using the FEI Vitrobot IV (Thermo Fisher Scientific). The vitrification was performed at room temperature at 100 % humidity. The data were collected on Titan Krios G2 (The Hormel Institute) operating at 300 kV equipped with Gatan K4 camera, using nominal magnification of x130,000 at a physical pixel size of 0.66 Å and total dose of 50 e^-^/Å. The data collection resulted in 15,970 and 18,149 movies for the complex and TfR1 alone, respectively.

### Image processing

The image processing for the icosahedral reconstruction was performed in CryoSPARC v4^37^. Particles were initially picked by “blob picker” and then automatically selected using “template picker.” Iterative rounds of 2D classification were performed to remove junk particles from further processing, and an ab-initio model was generated using C2 symmetry from the selected particle subset. After heterogenous refinement, 3D density was reconstructed using non-uniform refinement with global and local CTF refinement. Based on resolution improvement, local motion correction was performed for the VP1u- TfR1 complex data, whereas reference-based motion correction was performed for the TfR1 alone data. The motion corrected particles were re-refined, yielding 2.23 Å and 2.42 Å resolution for VP1u-TfR1 and TfR1-alone, respectively.

To resolve partial occupancy of VP1u on the dimeric TfR1, symmetry expansion followed by focused 3D classification was performed. The 681,957 particles refined with C2 symmetry were symmetry-expanded yielding 1,363,914 particles. Using a mask focused on one of the VP1u densities, 3D classification without alignment was performed to discard particles lacking VP1u. The same process was repeated for the other VP1u density on TfR1, yielding 1,017,894 particles. To account for the flexible interaction between VP1u and TfR1, the particles were subjected to flexible refinement, including 3D flex data prep, mesh prep, training, and reconstruction, yielding a final map at 2.41 Å resolution.

### Model building and refinement

The atomic model of the human TfR1^24^ was initially refined against the cryo-EM density of the TfR1-alone map by using ISOLDE^38^ and real-space refinement in PHENIX^39^. For the atomic model of VP1u, the density corresponding to VP1u was isolated from the VP1u-TfR1 complex map by applying a mask and, together with amino acid sequences of VP1u or VP1, was subjected to ModelAngelo^25^ in Relion 5.0 for *de novo* model building. The generated model for the VP1u RBD was refined independently by using ISOLDE^38^, combined with TfR1, and further refined against the VP1u-TfR1 complex map using ISOLDE^38^ and real-space refinement in PHENIX^39^. Final validation for the models was done using MolProbity^40^.

### Structural analysis and display

Contacts between VP1u RBD and TfR1 were identified as residues having atoms separated by less than 0.4-Å van der Waal’s radius in ChimeraX^41^. Map visualization and images were also generated in ChimeraX^41^.

### Quantification and statistical analysis

Statistical analyses were performed using GraphPad Prism version 10.3.1 (GraphPad Software, Boston, MA, USA). The specific statistical tests applied are indicated in the corresponding figure legends. Error bars represent standard deviation (SD), as specified in each legend. A p value < 0.05 was considered statistically significant in all analyses.

### Data availability

The cryo-EM maps and atomic coordinates of the TfR1 alone and the VP1u-TfR1 complex have been deposited in the EM database (https://www.emdatabank.org) and protein data bank (https://www.rcsb.org) under accession codes EMD-75944 (PDB entry 11QC) and EMD-75980 (PDB entry 11RN), respectively.

## Acknowledgments

We thank Dirk Grimm (University of Heidelberg, Germany) and Jean-Louis Reymond (University of Bern, Switzerland) for kindly providing the Jurkat and the HepG2 cells, respectively. This study was supported by a grant from the Swiss National Science Foundation (grant 320030_207850 to C.S.).

---

**Extended Data Fig. 1.**
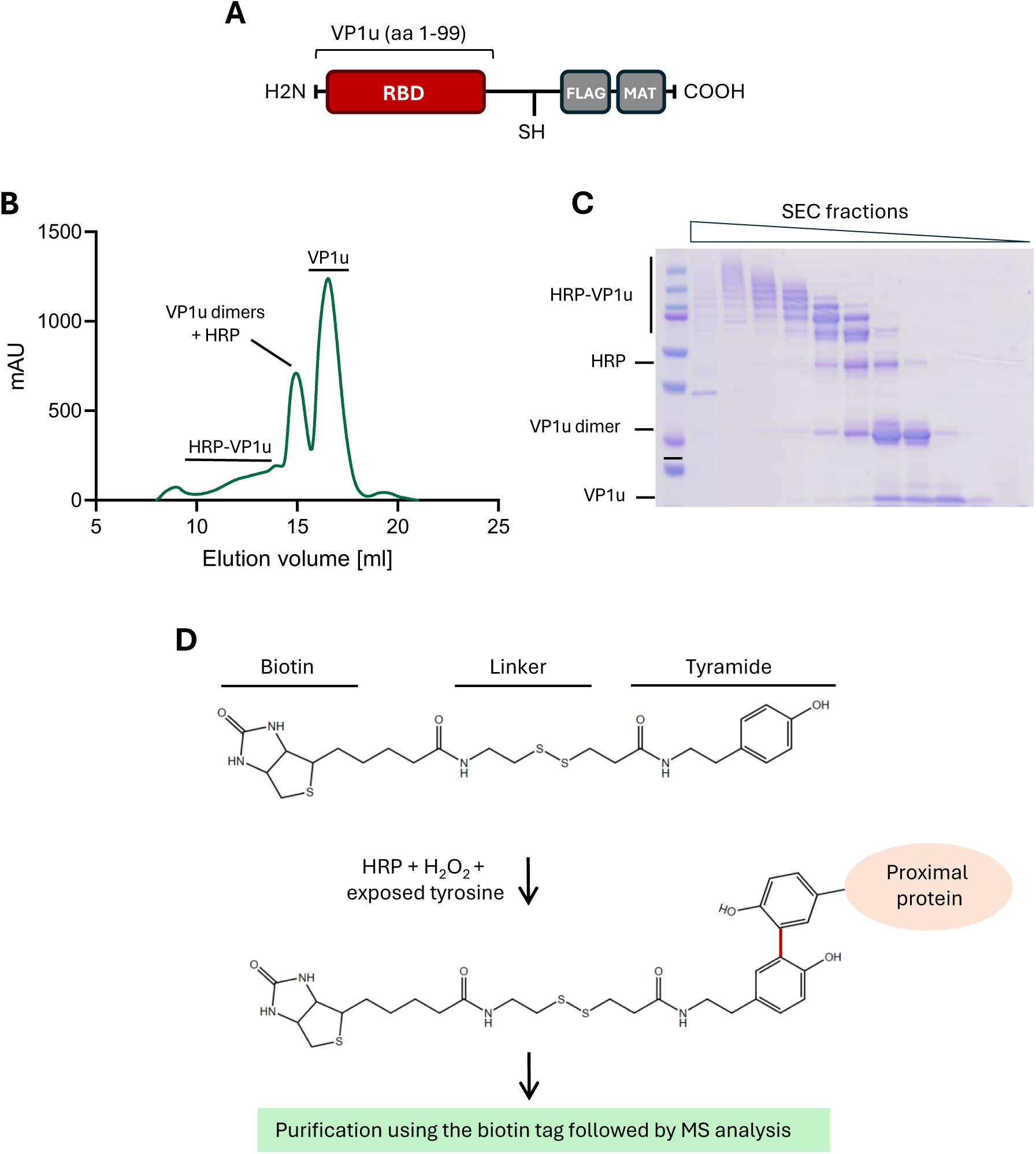
Schematic representation of the Selective Proteomic Proximity Labeling Assay Using Tyramide (SPPLAT). **(A)** Schematic depiction of the recombinant VP1u with a 128 amino acid truncation at its C-terminus. After conjugation of VP1u molecules to HRP, the reactants were separated using size-exclusion chromatography (SEC) **(B)** and collected fractions were analyzed by SDS-PAGE **(C)**. **(D)** Schematic illustration of the SPPLAT. HRP generates reactive tyramide radicals in the presence of H₂O₂, leading to covalent biotinylation of proteins in close proximity at the plasma membrane.

**Extended Data Fig. 2.**
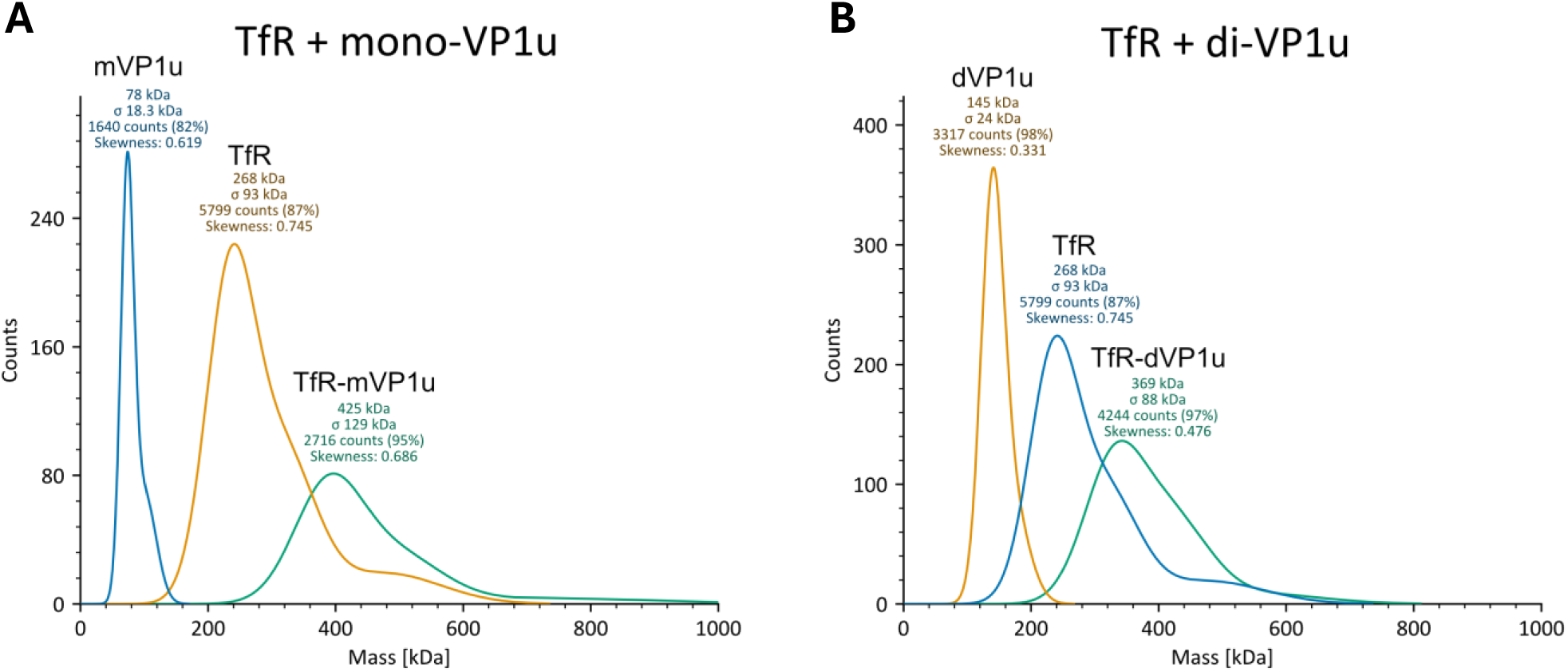
Mass photometry of the VP1u-TfR1 complexes. The mass photometry histograms of TfR, VP1u recombinants, and their mixtures show the molecular size distributions and the formation of TfR-VP1u complexes for the VP1u monomer **(A)** and VP1u dimer **(B)**.

**Extended Data Fig. 3.**
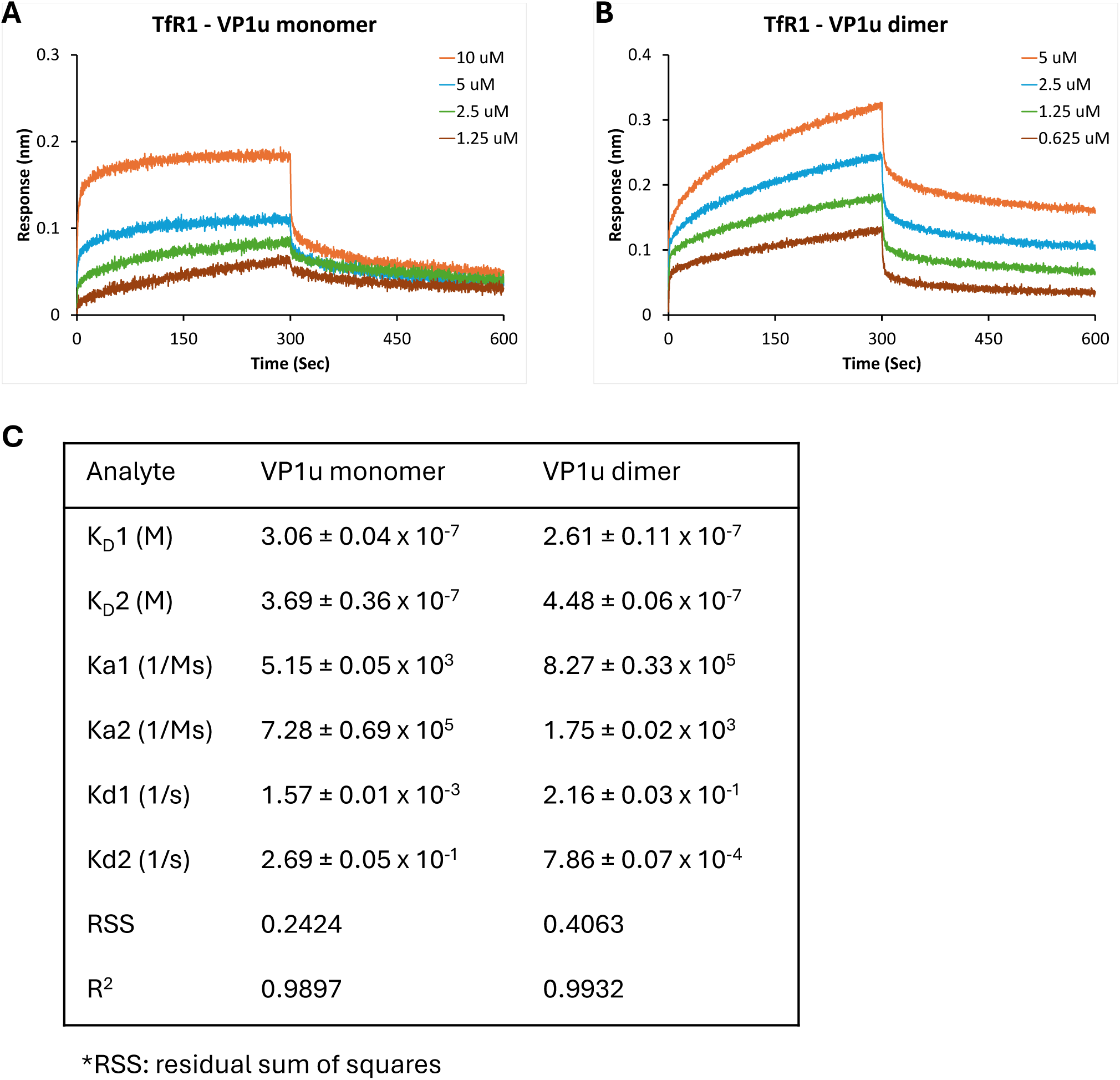
Binding kinetics of TfR1 with recombinant VP1u monomer and dimer. Binding of VP1u monomer **(A)** and dimer **(B)** to TfR1-Fc was measured by bio-layer interferometry using serial dilutions. TfR-Fc was immobilized on an anti-Fc biosensor surface. VP1u samples were incubated for 300 s, followed by 300 s dissociation phase. **(C)** Binding kinetics parameters are summarized in the table. Two binding modes, K_D_1 and K_D_2, were analyzed using a 2:1 heterogeneous ligand model.

**Extended Data Fig. 4.**
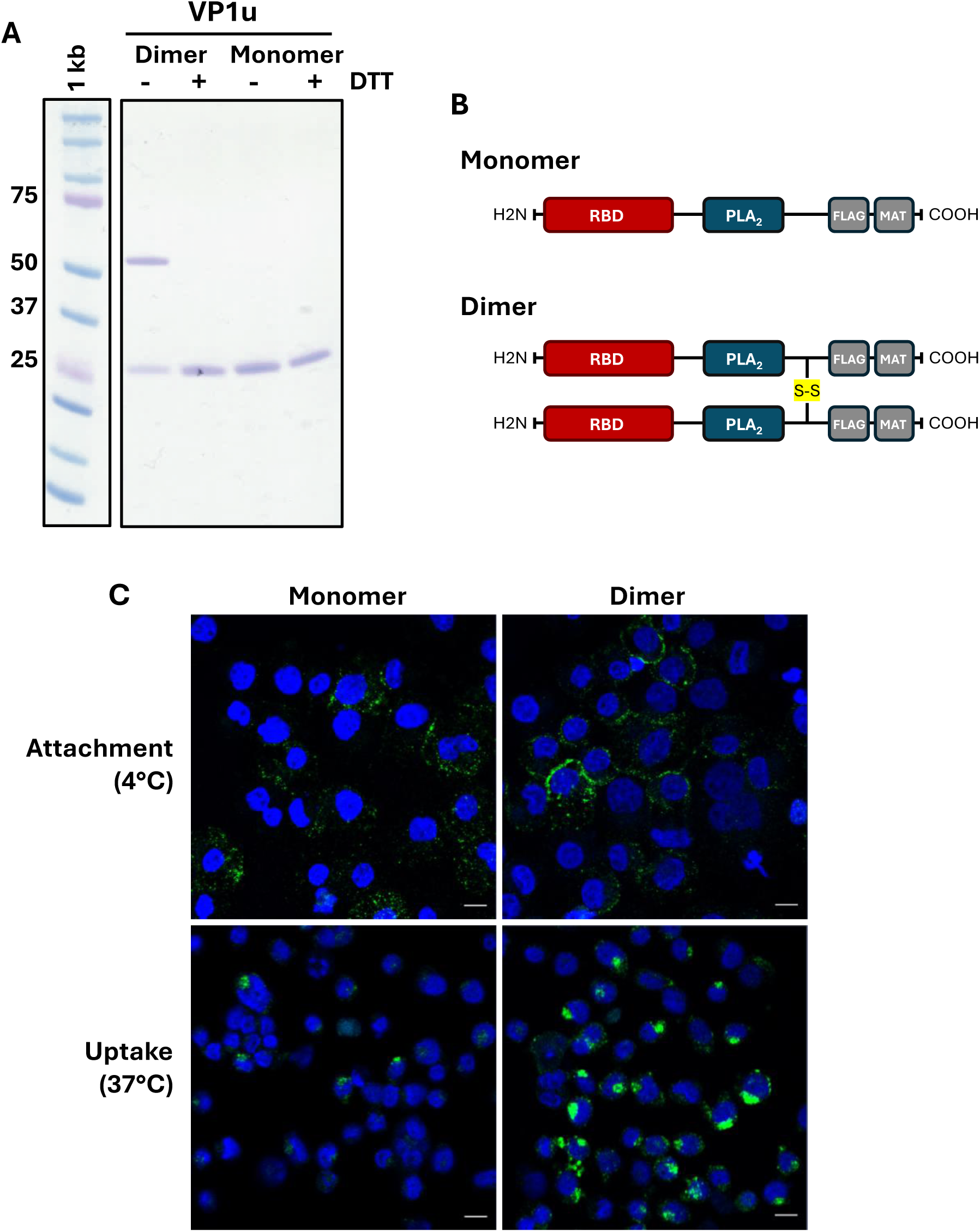
Binding and uptake of recombinant VP1u monomer and dimer in UT7/Epo cells. **(A)** SDS-PAGE analysis of purified recombinant VP1u proteins (monomer and dimer) used in this study. Molecular weight markers are shown on the left. Purified VP1u preparations were separated by electrophoresis and visualized by Coomassie staining. **(B)** Schematic representation of the VP1u monomer and dimer. The N-terminal receptor-binding domain (RBD) and the phospholipase A₂ (PLA₂) domain are indicated. Constructs contain C-terminal FLAG and MAT tags. One construct contains an engineered disulfide bond (S-S) to generate a covalent VP1u dimer. **(C)** Confocal microscopy images of UT7/Epo cells incubated with recombinant VP1u constructs. Cells were exposed to monomer or dimer VP1u proteins under conditions allowing cell surface binding (4 °C) and uptake (37 °C) and subsequently analyzed by confocal microscopy. Scale bar: 10 μm.

**Extended Data Fig. 5.**
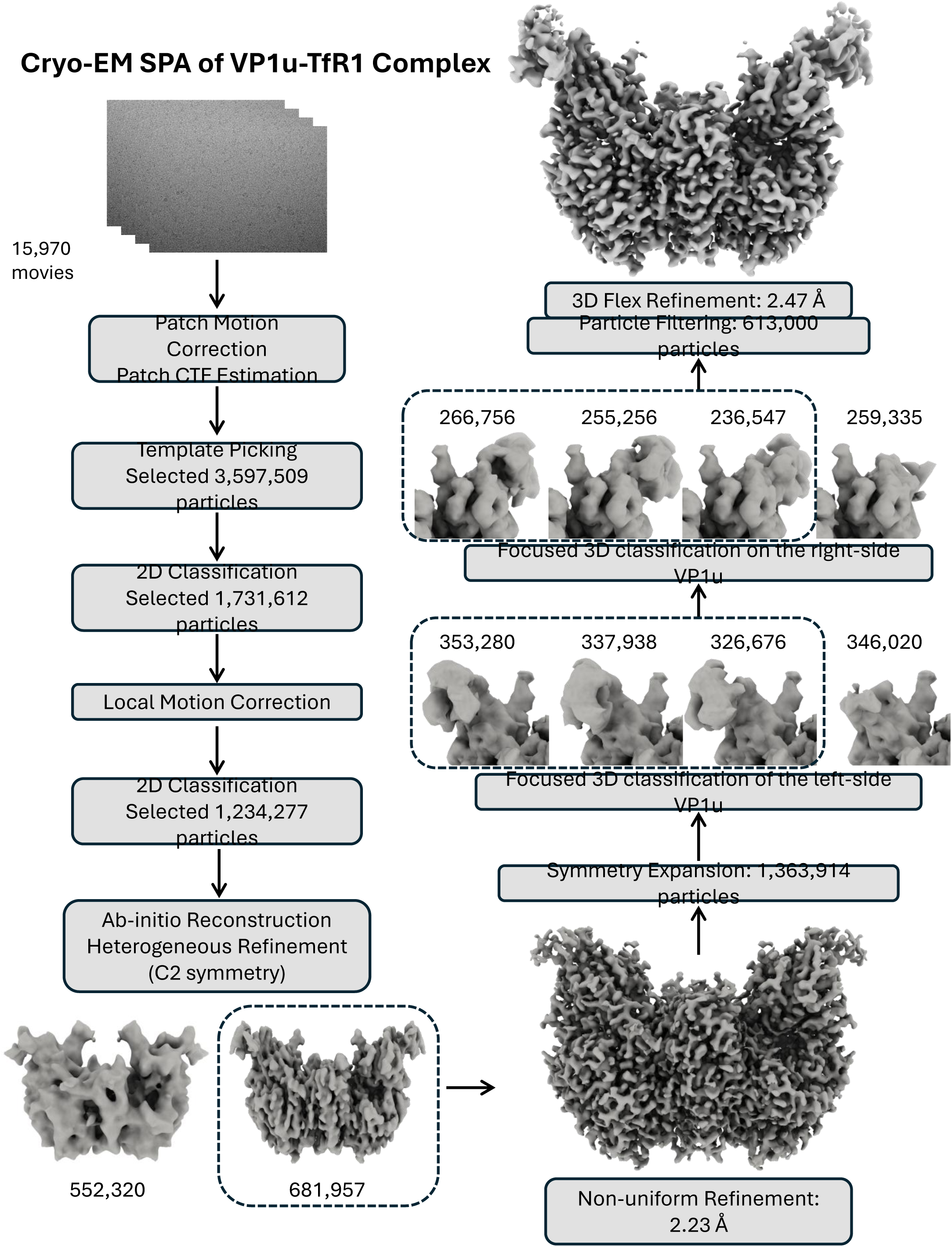
Schematic representation of the cryo-EM single particle analysis of VP1u-TfR1 complex.

**Extended Data Fig. 6.**
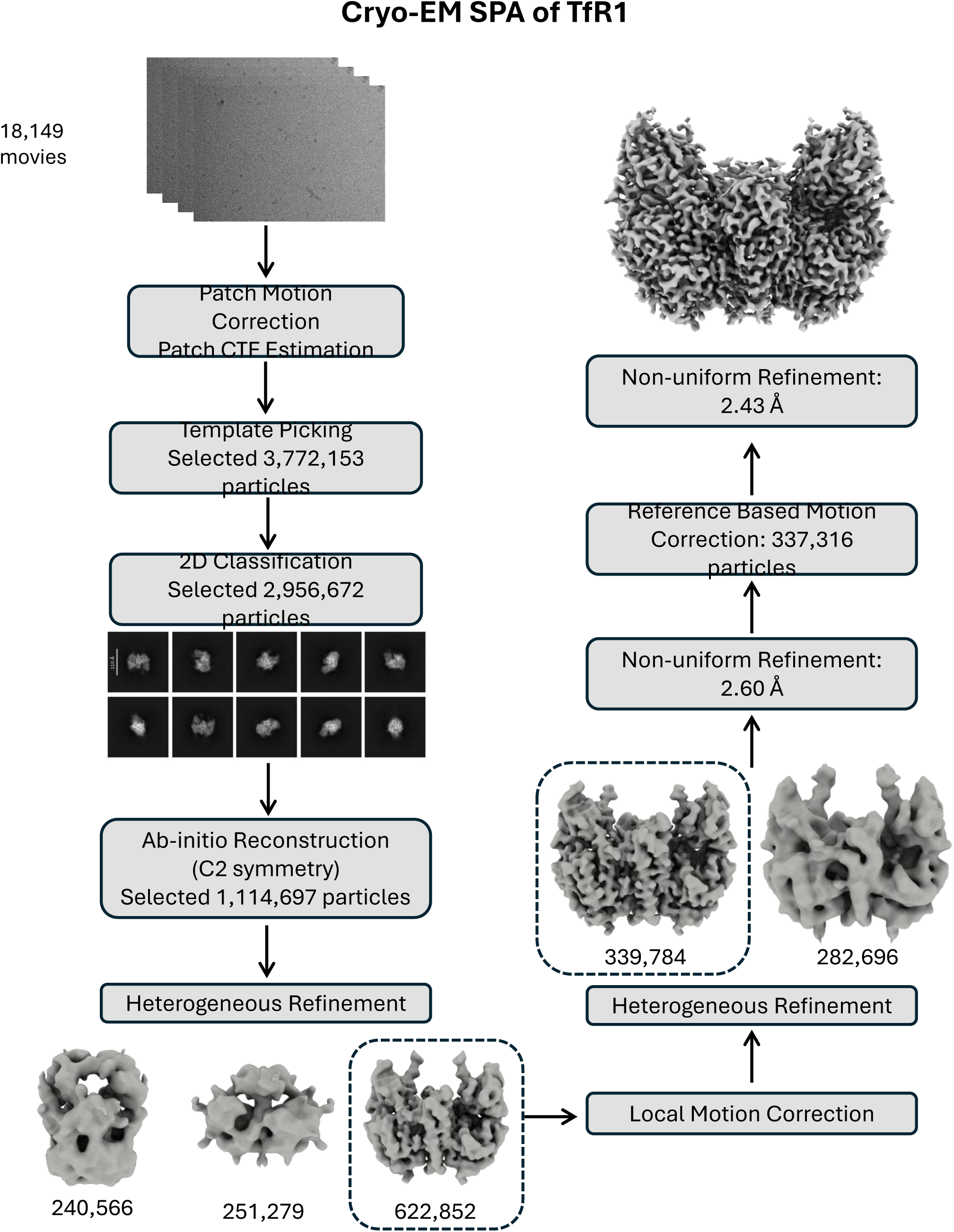
Schematic representation of the cryo-EM single particle analysis of TfR1.

**Extended Data Fig. 7.**
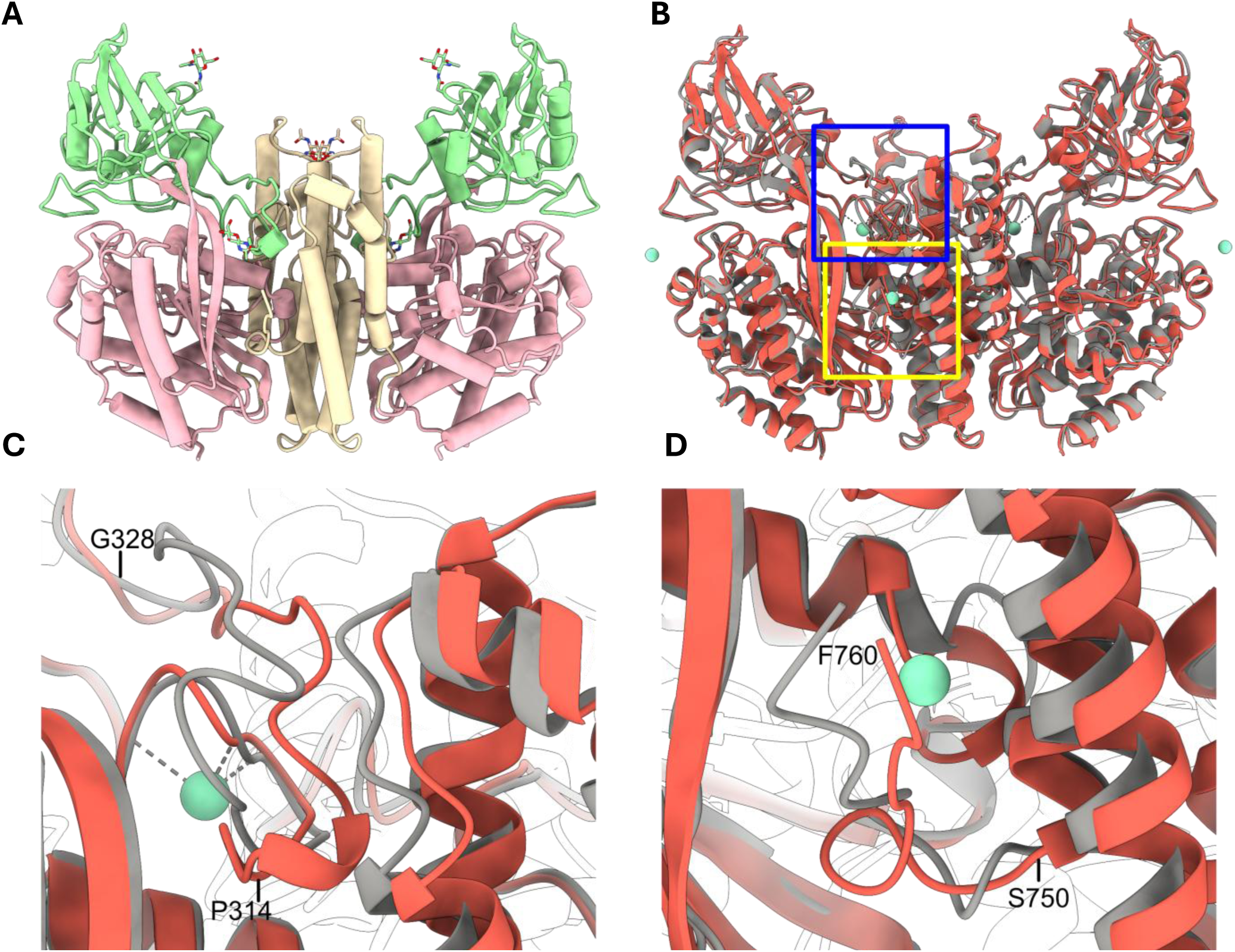
Comparisons of the TfR1 structures. **(A)** Atomic model of human TfR1 generated from the cryo-EM structure. Domains are colored as in Fig. 5. **(B-D)** The cryo-EM structure (red) aligned with human TfR1 crystal structure (gray). Sm^3+^ ions are shown in light green spheres. Regions highlighted by blue and yellow boxes are zoomed in panels C and D, respectively. **(C)** Conformational differences in the apical traverse loop (residues Pro 314–Gly 328) and adjacent residues of the helical domain. **(D)** Conformational variation in the C-terminal residues. **(E)** Comparison of the TfR1-alone structure (red) with the VP1u-TfR1 complex (blue). **(F)** Close-up view of the apical loop, showing the conformational change upon RBD binding (gray).

**Extended Data Table 1.**

| EMDB<br>PDB | TfR1<br>EMD-75944<br>11QC | VP1u-TfR1<br>EMD-75980<br>11RN |
| --- | --- | --- |
| **Data collection and processing** |  |  |
| Magnification | 130Kx | 130Kx |
| Voltage (kV) | 300 | 300 |
| Electron exposure (e<sup>-</sup>/Å<sup>2</sup>) | 50 | 50 |
| Defocus range (μm) | -1.0 to -2.0 | -1.0 to -2.0 |
| Pixel size (Å) | 0.66 | 0.66 |
| Initial micrographs (no.) | 18,145 | 15,696 |
| Final micrographs (no.) | 17,310 | 15,639 |
| Symmetry imposed | C2 | C2 (SymExp) |
| Particles (no.) | 337,316 | 613,000 |
| Map resolution (Å) | 2.43 | 2.41 |
| **Refinement** |  |  |
| Initial model used (PDB code) | 1CX8 | 11QC |
| Model resolution (Å) | 2.4 | 2.2 |
| FSC threshold | 0.143 | 0.143 |
| Model resolution range (Å) | 2.4 – 2.5 | 2.1 – 2.5 |
| Map sharpening B factor (Å<sup>2</sup>) | 103.0 | NA |
| Model composition |  |  |
| Non-hydrogen atoms | 10,198 | 11,202 |
| Protein residues | 1,278 | 1,398 |
| B factors (Å<sup>2</sup>) |  |  |
| Protein | 36.76 | 72.51 |
| R.m.s. deviations |  |  |
| Bond lengths (Å) | 0.003 | 0.003 |
| Bond angles (°) | 0.694 | 0.657 |
| Validation |  |  |
| MolProbity score | 0.94 | 1.02 |
| Clashscore | 0.79 | 1.04 |
| Poor rotamers (%) | 0.55 | 1.00 |
| Ramachandran plot |  |  |
| Favored (%) | 96.86 | 96.62 |
| Allowed (%) | 2.98 | 3.24 |
| Disallowed (%) | 0.16 | 0.14 |

